# Decoding subtype development and function in human pluripotent stem cell-derived midbrain dopaminergic neurons

**DOI:** 10.64898/2026.09.23.753784

**Authors:** Donghe Yang, Sejoon Choi, Jinghua Piao, Alessandro Evangelisti, Zhe Yang, Vittoria Dickinson Bocchi, Shkurte Ademi Donohue, Nidia Claros, Johannes Jungverdorben, Mega Sidharta, Youjun Wu, Hilda Amalia Pasolli, Tae Wan Kim, Ting Zhou, David Sulzer, Eugene V. Mosharov, Viviane Tabar, Lorenz Studer

**Author notes:** Co-Corresponding Authors: Lorenz Studer, Donghe Yang.

## Abstract

Midbrain dopaminergic (mDA) neurons comprise molecularly and functionally distinct subtypes with differential vulnerability in neurodegenerative and psychiatric disorders. However, the mechanisms specifying subtype identity remain poorly understood, and protocols for the selective derivation of human mDA subtypes are lacking.

Here we establish a strategy to derive substantia nigra (A9) and ventral tegmental area (A10) mDA neurons from human pluripotent stem cells (hPSCs). A9 identity is specified by dual-SMAD activation through Activin A and BMP7 at the midbrain floor-plate progenitor stage, whereas A10 identity is promoted by BMP inhibition. A9 mDA neurons are purified based on ALDH1A1 expression, and subtype identity is maintained by continued TGF-β modulation and ESRRB activation *in vitro* and upon transplantation *in vivo*. Single-cell RNA sequencing and biochemical analyses demonstrate that hPSC-derived A9 neurons exhibit increased oxidative phosphorylation, neuromelanin-like pigmentation, elevated dopamine synthesis and release, and electrophysiological properties characteristic of A9 mDA neurons *in vivo*. Integration with human fetal midbrain datasets confirms strong transcriptional concordance between *in vitro*-derived and *in vivo* mDA subtypes.

Neuromelanin-like structures produced by hPSC-derived A9 neurons trigger pro-inflammatory cytokine secretion from hPSC-derived microglia, and A9 neurons are primed to upregulate MHC-I genes in response to interferon-γ, features which may contribute to the selective vulnerability of A9 neurons. Together, these results establish a robust *in vitro* platform to interrogate human mDA subtype development, function, and selective vulnerability, enabling mechanistic studies relevant to Parkinson’s disease and the development of cell-based therapies.

## Introduction

The midbrain dopaminergic (mDA) system regulates voluntary movement, reward processing and cognition, and its dysfunction contributes to both neurodegenerative and psychiatric disorders. A9 neurons in the substantia nigra pars compacta (SNpc) and A10 neurons in the ventral tegmental area (VTA) are distinguished by their anatomy, molecular identity and physiology^1–3^. A9 neurons innervate the dorsolateral striatum via the nigrostriatal pathway and are essential for motor control. In Parkinson’s disease (PD), A9 mDA neurons are selectively and progressively lost, whereas neighboring A10 neurons are relatively spared^4^. Conversely, A10 neurons participate in meso-corticolimbic circuits implicated in depression, addiction and schizophrenia^5^. Despite extensive efforts to derive human mDA neurons for disease modeling and cell-based therapies^6–8^, including ongoing clinical trials in patients with advanced Parkinson’s disease^9,10^, the developmental programs and regulatory networks that establish and maintain human A9 versus A10 identity, and that underlie their selective vulnerability, remain poorly defined.

A major barrier to studying human A9 and A10 mDA subtypes is limited access to these neurons: they are rare, anatomically inaccessible, difficult to isolate and are consistently underrepresented in single cell studies. Atlases of the developing and adult midbrain have identified a set of subtype-specific markers, including *LMO3, SOX6* and *ALDH1A1* for A9 neurons and *CALB1/CALB2* and *OTX2* expression for A10 lineage, and have linked these molecular states to differential vulnerability in PD^11–15^. However, marker catalogs alone do not reveal the upstream cues that specify these fates, or the mechanisms required to stably maintain subtype identity, which are prerequisites for generating enriched, well-defined human mDA subtypes from hPSCs.

Protocols to derive mDA neurons from hPSCs yield mixed populations whose subtype identity is variable and often drifts with maturation or upon transplantation *in vivo*^8,16–18^. Prior efforts to bias subtype outcome have focused on WNT pathway tuning, SMAD inhibition timing or transcription factor (*ASCL1* or *OTX*2) overexpression^19–22^. However, most of those studies yield inconsistent effects as illustrated by the lack of hallmark A9 features such as *ALDH1A1* expression, and did not provide definitive protocol for selective generation of A9 or A10 neurons that can be used for modeling basal ganglia circuitry. One recent study used a genetic reporter strategy to successfully isolate A10 lineage within a mixed population of hPSC-derived mDA neurons, but the gene regulatory networks driving A10 specification remain unclear^5^. Furthermore, the field lacks prospective, reporter-free strategies to purify and longitudinally track subtype-committed progenitors and neurons.

Here we establish a broadly applicable, two-stage framework to generate human A9- and A10-like mDA neurons with subtype-resolved transcriptional, electrophysiological, metabolic and neuroimmune phenotypes. First, we show that transient exposure of ventral midbrain progenitors to Activin A and BMP7 biases specification toward an *ALDH1A1⁺* A9 program, whereas BMP pathway inhibition drives *CALB1⁺* A10 fate. Second, guided by multiome sequencing-derived regulatory predictions, we identify a postmitotic “A9 maintenance” condition which involves co-activation of TGF-β/BMP and ESRRB signaling that stabilizes A9 hallmark features at later stages that otherwise diminish *in vitro*.

Beyond protocol development, we introduce practical, reporter-free strategies to isolate and track subtype trajectories over time by combining surface-marker gating (CD56/CD171 and CD99) with ALDH enzymatic activity. This enables prospective purification of A9- and A10-committed progenitors and neurons for downstream molecular and functional analyses. Functionally, hPSC-derived A9-like neurons exhibit higher oxidative phosphorylation capacity, ultrastructural mitochondrial features consistent with elevated respiratory demand, and selective neuromelanin-like pigment formation, whereas A10-like neurons show signatures consistent with greater excitatory synaptic input, in line with their established circuit roles. Finally, we leverage this platform to model subtype-relevant neuroimmune interactions: neuromelanin-like material from A9 cultures activates hPSC-derived microglia to secrete pro-inflammatory cytokines, and A9-like neurons mount a stronger MHC-I response to inflammatory stimulation than A10-like neurons, supporting a tractable *in vitro* system to study immune-mediated contributions to selective A9 vulnerability.

Together, our work provides a mechanistically grounded roadmap for subtype specification and maintenance of human mDA neurons, enabling scalable production, purification and functional interrogation of A9- and A10-like populations for PD pathobiology, psychiatric circuit modeling and the development of subtype-appropriate cell replacement strategies.

## Results

### Identification of potential time windows and regulators of mDA subtype specification

To define the *in vitro* time window and candidate regulators underlying mDA subtype differentiation, we reanalyzed our previously published single-cell RNA-seq (scRNA-seq) datasets profiling hPSC-derived mDA populations across developmental stages (days 16, 25, and 40) (Extended Figs. 1a,b)^8^. This reanalysis resolved discrete populations, including ventral midbrain progenitors (*WNT1*⁺*SHH*⁺), neuroblasts (*NR4A2*⁺*LMX1A*⁺*FOXA2*⁺), and emerging putative mDA neuron subtypes marked by high *LMO3* expression (A9-like) or *CALB2* expression (A10-like) (Extended Fig. 1c).

To investigate the diversification of A9- and A10-like subtypes and their developmental trajectories, we performed *in silico* pseudotime reconstruction using the URD algorithm, designating MKI67⁺ proliferating progenitors as the developmental origin (Extended Fig. 1d)^23^. The URD analysis suggested that the divergence of A9- and A10-like subtypes is established by day 25 *in vitro*, corresponding to the early neuronal stage when tyrosine hydroxylase (*TH*), a functional mDA marker^24^, begins to be expressed. Notably, trajectories 1 and 3 likely represent the developmental paths of A9 and A10 lineages, respectively, as indicated by the differential expression of subtype-specific markers (Extended Figs. 1e,f).

These computational findings are further supported by immunofluorescence (IF) analyses of day 25 and day 30 hPSC-mDA populations. A subset of TH⁺ cells at these stages expressed ALDH1A1, a well-characterized A9 marker^1,25^, consistent with the hypothesis that subtype specification occurs prior to the appearance of TH⁺ mDA neurons (Extended Fig. 1g).

To identify signaling pathways potentially involved in early subtype specification, we focused on neuronal progenitors present before day 25. Cell-cell communication analysis using CellChat revealed that BMP and TGF-β signaling pathways were highly enriched in the progenitor populations (Extended Figs. 1h,i)^26^, implicating them in the regulation of mDA subtype development.

### BMP7 and Activin A promote specification of the A9 mDA subtype

To define signaling requirements for mDA subtype specification (Fig. 1a), we generated a CRISPR–Cas9 knock-in *PITX3*::GFP reporter line in the H9 background, as *PITX3* marks postmitotic mDA neurons and is required for A9 development^27,28^. The resulting line was validated by CRISPRa and efficiently differentiated into PITX3⁺TH⁺ mDA neurons using our established “Boost+” protocol^8^ (Fig. 1b). To further assess the dynamics of mDA differentiation, we incorporated additional surface markers: CD99, which labels neuroblasts or progenitors, and CD171 or CD56, which mark differentiated neurons^29,30^. This combination enabled tracking of mDA progression. Using these markers, we performed flow cytometry (FACS) analysis and identified that exposure to Activin A (10 ng/mL) from day 17 to day 25 significantly increased the proportion of cells expressing PITX3 within both the CD99⁺ and CD171⁺ populations (Fig. 1c and Extended Fig. 2a). Conversely, inhibition of Activin signaling with the small molecule inhibitor SB431542 (SB) resulted in a reduction in the proportion of PITX3⁺ mDA neurons (Extended Fig. 2b).

**Fig. 1.**
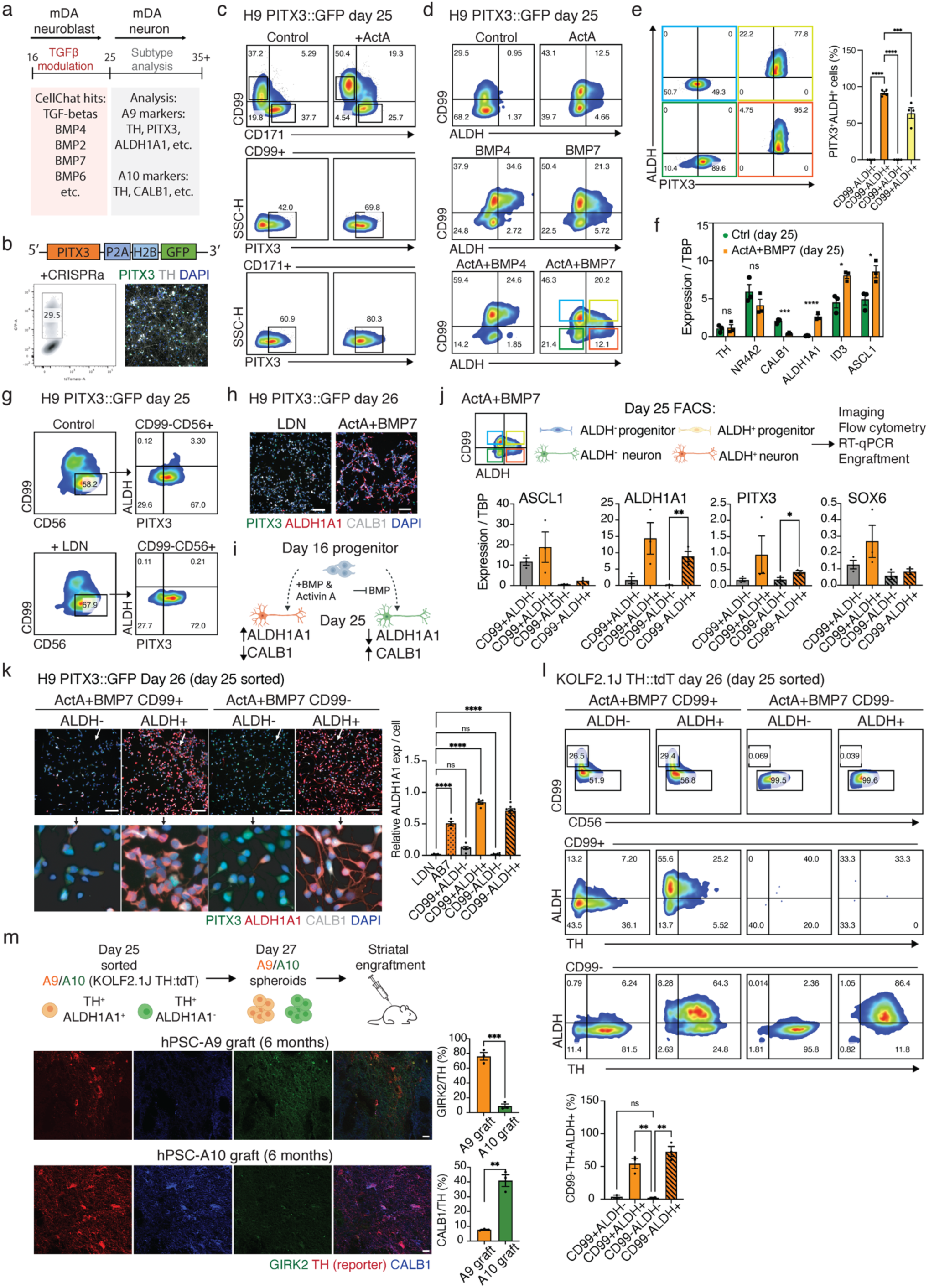
BMP7 and Activin A promote specification of the A9 mDA subtype. **a**, Experimental schematic of TGFβ pathway modulation during mDA progenitor patterning. **b,** PITX3::GFP reporter design (top), flow cytometry of PITX3 CRISPRa activation (middle), and immunofluorescence (IF) for PITX3 and TH at D30 (bottom). **c,** Flow cytometry of PITX3, CD171 and CD99 at D25. **d,** Flow cytometry of ALDH activity and CD99 at D25. **e,** Flow cytometry of PITX3 expression and ALDH activity in the indicated gated subpopulations, with quantification of the percentage of PITX3+ALDH+ cells in each subpopulation. One-way ANOVA; *n* = 4. **f,** RT–qPCR of the indicated genes in D25 control and Activin A/BMP7-patterned populations. Student’s *t*-test; *n* = 3. **g,** Flow cytometry of PITX3 and ALDH activity in the indicated CD99-CD56+ populations. **h,** IF for PITX3, ALDH1A1 and CALB1 at D26. Scale bars, 100 μm. **i,** Schematic illustrating mDA subtype patterning through modulation of TGF-β signaling. **j,** Schematic of the day 25 sorting strategy and RT–qPCR analysis of the indicated genes in sorted day 25 populations. One-way ANOVA; *n* = 3–4. **k,** IF for ALDH1A1, PITX3 and CALB1 at D26 in populations derived from D25 sorted cells (left), with quantification of relative ALDH1A1 expression (right). Scale bars, 100 μm. One-way ANOVA; *n* = 4–8. **l,** Flow cytometry of TH and ALDH activity in the indicated CD99− and CD99+ populations, with quantification of the percentage of CD99-TH+ALDH+ cells. One-way ANOVA; *n* = 3. **m**. Schematic of hPSC-A9/A10 transplantation into the mouse striatum (top); IF staining for GIRK2, TH and CALB1 in A9 and A10 grafts at 6 months post-transplantation, with quantification of GIRK2+TH+ and CALB1+TH+ cells in the grafts. Student’s *t*-test; *n* = 3. All error bars in Fig. 1 represent s.e.m.

We next combined the *PITX3*::GFP reporter with the AldeRed assay, which detects cellular aldehyde dehydrogenase (ALDH) activity ^25,31,32^ and, in our system, serves as a readout of ALDH1A1 expression. This established a live-cell FACS assay to quantify and isolate PITX3⁺ALDH⁺ A9 mDA neurons from hPSC-derived cultures (Figs. 1d,e). Using this assay, we evaluated the effects of various BMP ligands alongside Activin A and found that BMP7 (20 ng/mL), in combination with Activin A, significantly enhanced the generation of CD99⁻ALDH⁺PITX3⁺ neurons (Figs. 1d,e and Extended Fig. 2c). RT-qPCR confirmed increased *ALDH1A1* and reduced *CALB1*, an A10 marker (Fig. 1f and Extended Fig. 2d). Using the KOLF2.1J TH::tdTomato reporter line^33^, we demonstrated that inhibition of Activin and BMP signaling pathways with SB and LDN193189 (LDN), respectively, abolished the generation of ALDH⁺TH⁺ A9 mDA neurons (Extended Fig. 2e).

Notably, BMP4 treatment (20 ng/mL) failed to generate the desired CD99⁻ALDH⁺ A9 population (Fig. 1d and Extended Fig. 2c). Consistently, day 25 cultures patterned with Activin A and BMP7 yielded neurons co-expressing ALDH1A1 and TH, whereas Activin A and BMP4 did not produce a comparable TH⁺ population (Extended Fig. 2f). scRNA-seq further showed that Activin A/BMP7-patterned cells were transcriptionally similar to control mDA neurons and enriched for genes associated with synaptic function and ion transport (Extended Figs. 2g–i). By contrast, Activin A/BMP4-patterned cells adopted a distinct profile enriched for epithelial differentiation genes, indicating divergence from the mDA lineage (Extended Fig. 2i). Together, these data identify BMP7 as a key regulator of A9 mDA neuron specification, consistent with *in vivo* evidence that SMAD1 inactivation impairs the generation of *TH*⁺*SOX6*⁺ substantia nigra neurons^34^.

Because BMP ligand treatment reduced expression of the A10 marker *CALB1* (Extended Fig. 2d), we next tested whether BMP inhibition would instead favor A10 identity. Treatment with the BMP inhibitor LDN from days 17 to 25 reduced ALDH1A1 and increased CALB1 expression at days 25 and 30 (Figs. 1g,h and Extended Figs. 2j–l). Thus, Activin A/BMP7 biases differentiation toward an A9 fate, whereas BMP inhibition favors A10 specification (Fig. 1i).

### Purification of mDA neuron subtypes

We then asked whether A9 progenitors and neurons could be prospectively isolated from A9-enriched cultures using CD99 and ALDH activity. RT-qPCR confirmed that sorted ALDH⁺ neurons expressed higher *ALDH1A1* and *PITX3* than ALDH⁻ cells (Fig. 1j), and overnight replating showed enrichment of ALDH1A1⁺ neurons by IF (Fig. 1k).

This strategy also enabled longitudinal tracking of A9 differentiation. Using the KOLF2.1J TH::tdTomato line, we isolated CD99⁺ALDH⁺ progenitors at day 25. Within 24 h, these cells rapidly transitioned toward a neuronal fate, marked by CD99 downregulation and TH upregulation, while retaining ALDH1A1 expression (Fig. 1l). These data reveal a CD99⁺ALDH1A1⁺TH⁻ progenitor intermediate that precedes A9 mDA neuron differentiation, indicating that subtype commitment can occur before TH onset. However, because CD99⁺ALDH⁺ cells also generated TTR⁺FOXA2^low^TH^low^ non-mDA/choroid plexus-like contaminants, subsequent analyses focused on purified CD99⁻ALDH⁺ A9-like neurons (Extended Fig. 2m)^35,36^.

We next tested protocol generalizability in additional hPSC/iPSC lines, including MSK-SRF001 and J1^37,38^. BMP inhibition for A10 specification and Activin A/BMP7 activation for A9 specification reproducibly generated the corresponding mDA subtypes across lines, supporting the robustness and broader applicability of this strategy (Extended Fig. 2n).

Finally, we asked whether purified A9- and A10-like cells could engraft *in vivo* while retaining subtype-specific features. Day 25 purified A9 (TH⁺ALDH⁺) and A10 (TH⁺ALDH⁻) populations derived from the KOLF2.1J TH::tdTomato line were reaggregated for 2 days, transplanted into the striatum of NSG mice and analyzed 6 months later. Grafted neurons survived in host tissue and maintained subtype-biased marker expression: A9 grafts contained a higher fraction of GIRK2⁺TH::tdTomato⁺ neurons than A10 grafts (76% versus 9%), whereas A10 grafts contained more CALB1⁺TH::tdTomato⁺ neurons than A9 grafts (41% versus 8%) (Fig. 1m). Together, these findings establish a robust, prospectively isolatable and transplantable human platform for generating molecularly distinct A9- and A10-like mDA lineages, enabling subtype-resolved studies of human dopaminergic development and therapeutic potential.

### Single-cell multiomic profiling of midbrain dopaminergic subtypes

Following mDA subtype patterning, we performed scRNA-seq and single-cell multiome profiling of day 25 A9- and A10-patterned cultures to define the molecular basis of subtype segregation. Cell populations were annotated using *CD99*, CD56/*NCAM* and CD171/*L1CAM*, together with markers of midbrain floor plate progenitors (*FOXA1*, *FOXA2*, *OTX2*, *LMX1A* and *LMX1B*), mDA neurons (*TH*, *DDC*, *EN1*, *PITX3* and *NR4A2*) and A9/A10 identity (*ALDH1A1* and *SOX6* for A9; *CALB1* and *CALB2* for A10) (Figs. 2a,b and Extended Figs. 3a–c)^8^. This analysis identified distinct *ALDH1A1*- and *CALB1*-expressing populations across neuroblast, early DA neuron and differentiated DA neuron states (Figs. 2a,b).

**Fig. 2.**
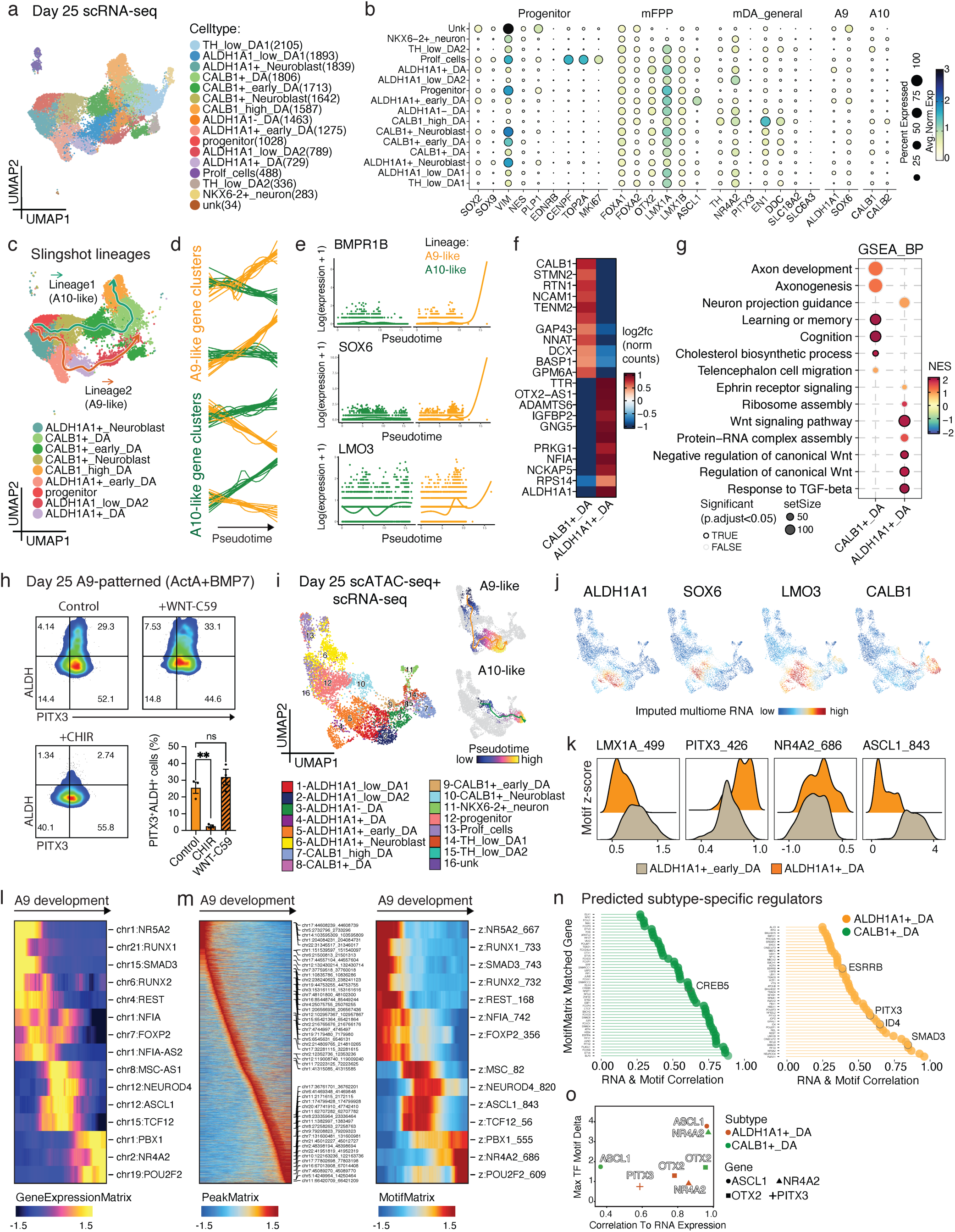
Single-cell multiomic profiling of midbrain dopaminergic subtypes. **a**, UMAP of scRNA-seq profiles from D25 A9- and A10-patterned mDA populations. **b,** Dot plot showing expression of the indicated gene categories across annotated clusters. **c,** Slingshot lineage inference showing two trajectories corresponding to A10-like (lineage 1) and A9-like (lineage 2) development. **d,e,** Gene-module scores (d) and A9 marker expression (e) plotted along pseudotime. **f,g,** Heatmap of differentially expressed genes (f) and pathway enrichment analysis (g) for CALB1+ DA and ALDH1A1+ DA clusters. **h,** Flow cytometry of PITX3 expression and ALDH activity in D25 A9-patterned populations under WNT modulation, with quantification of PITX3+ALDH+ proportions. One-way ANOVA; *n* = 3; error bars, s.e.m. **i,** UMAP of D25 single-cell Multiome (RNA+ATAC) profiles from A9- and A10-patterned populations with cell-type annotations (left) and pseudotime trajectories for A9- and A10-like development (right). **j,** UMAP feature plots showing imputed expression of the indicated genes. **k,** Z-scored enrichment of transcription factor motifs associated with dopaminergic genes in ALDH1A1+ clusters. **l,** Gene expression dynamics along the A9-like pseudotime trajectory. **m,** Accessibility of selected ATAC-seq peaks (left) and associated motif enrichments (right) along A9-like pseudotime. **n,** Candidate A9 and A10 regulators predicted by integrating motif enrichment with RNA expression. **o,** Selected predicted regulators highlighted in (n).

To reconstruct subtype-specific differentiation paths, we performed Slingshot analysis on *ALDH1A1*⁺ and *CALB1*⁺ clusters, using the progenitor cluster as the developmental origin^39^. The inferred trajectories resolved A10-like and A9-like lineages, corresponding to trajectories 1 and 2, respectively (Fig. 2c). TradeSeq analysis identified A9- and A10-specific transcriptional programs with opposing pseudotime dynamics (Fig. 2d)^40^. Consistent with lineage divergence, *SOX6* and *LMO3* increased along the A9 trajectory, as did *BMPR1B*, in agreement with BMP7-mediated A9 patterning (Fig. 2e).

We next compared differentiated *ALDH1A1*⁺ and *CALB1*⁺ DA neuron clusters by differential expression and GSEA. *ALDH1A1* was enriched in A9 DA neurons, whereas *CALB1* was enriched in A10 DA neurons (Fig. 2f). A10 neurons showed enrichment of gene programs associated with cognition and learning, consistent with ventral tegmental area dopaminergic neuron function. By contrast, A9 neurons were enriched for TGF-β signaling and negative regulation of Wnt signaling, matching the signaling conditions used for A9 patterning (Fig. 2g). To test the role of Wnt signaling, we treated Activin A/BMP7-patterned cultures with either the Wnt activator CHIR99021 (2 μM) or the Wnt inhibitor WNT-C59 (2 μM) from days 17 to 25. FACS analysis at day 25 showed that Wnt activation reduced the proportion of ALDH1A1⁺PITX3⁺ cells, indicating impaired A9 specification (Fig. 2h), consistent with prior evidence that excessive Wnt signaling disrupts SNc dopaminergic neuron development^22,41^.

We also examined MEK signaling, given prior evidence that MEK inhibition accelerates neuronal differentiation^42,43^. Treatment with the MEK inhibitor (MI) PD0325901 (10 μM) from days 17 to 25 increased expression of genes associated with neuronal maturation, synaptic function and neurotransmitter transport by GSEA (Extended Figs. 3d,e). In A9-patterned cultures, MI reduced CD99⁺ cells and increased CD56⁺ cells at day 25 (Extended Figs. 3f–h). This effect was most pronounced in the CD56⁻CD99⁺ALDH⁺ population, whereas the CD99⁻CD56⁺ALDH⁺ population was largely unchanged (Extended Figs. 3i,j), suggesting that MI primarily accelerates the transition from progenitor-like to neuronal states.

To further define the regulatory landscape of subtype specification, we performed single-cell multiome profiling of A9- and A10-enriched cultures (Fig. 2i and Extended Fig. 4a). Multiome cells were annotated by label transfer in ArchR using the day 25 scRNA-seq dataset as reference^44^. After RNA imputation with MAGIC^45^, A9 markers *ALDH1A1*, *SOX6* and *LMO3*, and the A10 marker *CALB1*, showed distinct expression patterns, confirming the presence of A9- and A10-like lineages in the multiome dataset (Fig. 2j and Extended Fig. 4b).

Chromatin accessibility analysis of *ALDH1A1*^+^ clusters revealed stage-specific regulatory dynamics during mDA differentiation. Motifs for *LMX1A* and *ASCL1*, associated with midbrain progenitor identity, were enriched in early neurons, whereas motifs for *PITX3* and *NR4A2*, associated with postmitotic DA neuron maturation, were more accessible in differentiated neurons (Fig. 2k). Comparison of differentially accessible peaks between *ALDH1A1*⁺ and *CALB1*⁺ early DA clusters revealed enrichment of *ID* family motifs in *ALDH1A1*⁺ cells and *FOXA* motifs in *CALB1*⁺ cells, with stronger *ID4* footprinting in the *ALDH1A1*⁺ early DA cluster (Extended Figs. 4c–e). These findings suggest that A9 and A10 lineages acquire distinct regulatory states early during subtype specification.

Finally, we integrated RNA expression with motif accessibility at differentially accessible peaks to nominate candidate regulators of A9 and A10 development (Figs. 2l,m). This analysis identified subtype-biased regulatory programs, including *CREB5* as a candidate A10 regulator and *SMAD3*, *PITX3* and *ESRRB* as candidate positive regulators of A9 development (Figs. 2n,o). Although *OTX2* was detectable in both subtypes at day 25, its RNA–motif correlation and maximum motif delta were higher in *CALB1*⁺ DA cells, consistent with prior reports linking *OTX2* to A10 identity^14^. Together, these analyses define coordinated transcriptomic and chromatin-based programs underlying human A9/A10 lineage segregation and nominate candidate regulators for mechanistic dissection of mDA subtype specification.

### Activin A, BMP4, and ESRRB agonist cooperatively maintain mature A9 mDA neuron characteristics

Although A9-enriched cultures showed robust *ALDH1A1* expression by day 25, this marker declined during later differentiation (Fig. 3a and Extended Fig. 5a). Because *ALDH1A1* is maintained in A9 neurons *in vivo*, we sought culture conditions that preserve A9 identity after subtype patterning. We therefore tested Activin A, BMP4 and BMP7, signals implicated in *ALDH1A1* induction during progenitor patterning, alone or in combination during postmitotic maturation (Fig. 3b).

**Fig. 3.**
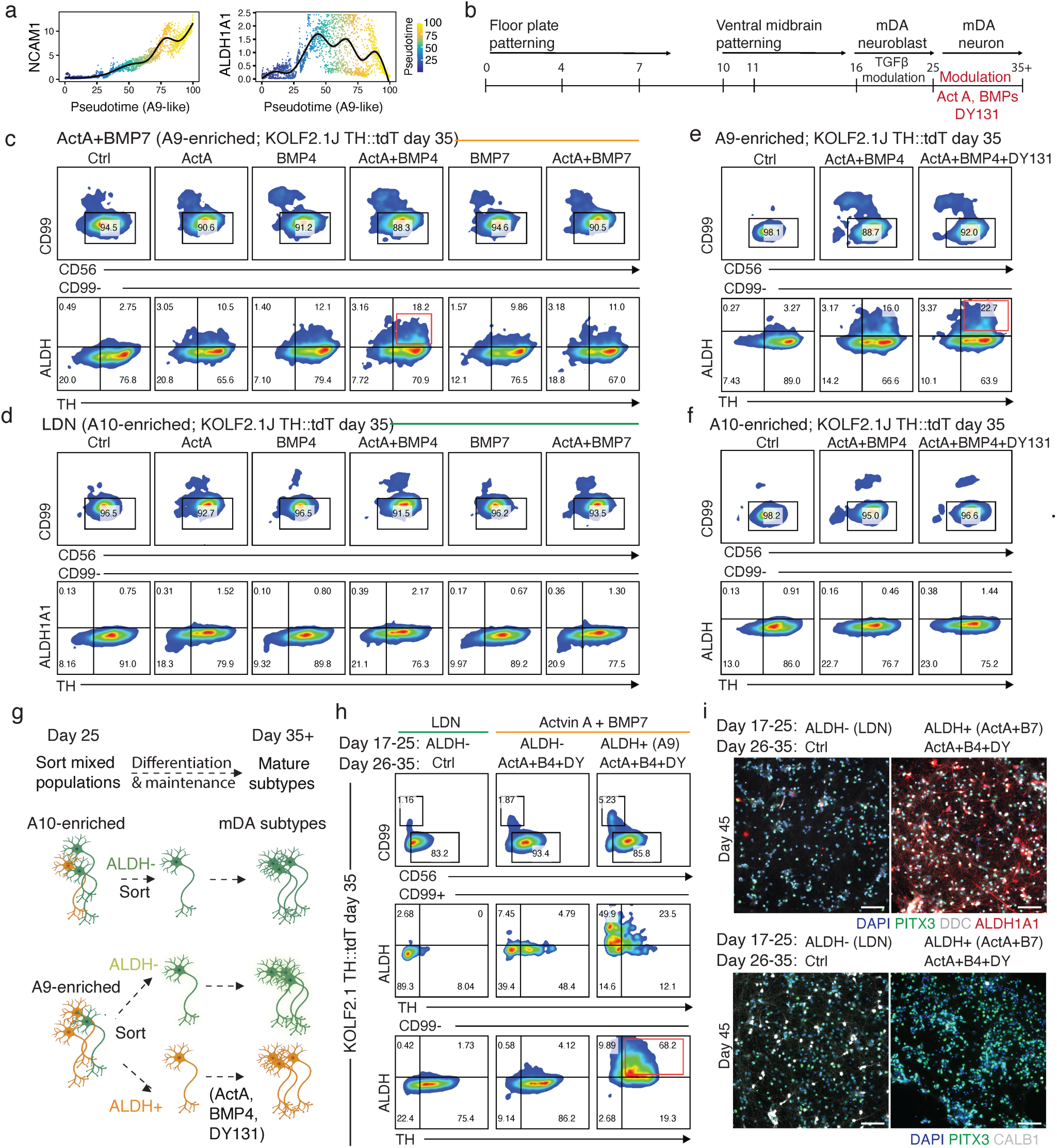
Activin A, BMP4 and an ESRRB agonist cooperatively maintain mature A9 mDA neuron characteristics. **a**, Expression of *NCAM1* and *ALDH1A1* along the A9-like pseudotime trajectory. **b,** Schematic of the experimental design and timing of pathway modulation. **c,d,** Representative flow cytometry of TH expression and ALDH activity within gated CD99-populations generated under the indicated conditions in D35 A9-patterned (c) and A10-patterned (d) cultures. **e,f,** Representative flow cytometry of TH expression and ALDH activity within gated CD99-populations generated under the indicated A9-maintenance conditions in D35 A9-patterned (e) and A10-patterned (f) cultures. **g.** Schematic of D25 sorting of A9- and A10-patterned populations and strategies to assess the effects of A9-maintenance conditions. **h,** Flow cytometry of TH expression and ALDH activity in CD99− and CD99+ fractions at D35 derived from D25 FACS-sorted subpopulations cultured under the indicated conditions. **i,** Representative IF images of D45 populations stained for PITX3, DDC and ALDH1A1 (top) or PITX3 and CALB1 (bottom) under the indicated conditions. Scale bars, 100 µm.

FACS and IF analyses at day 35 showed that Activin A and BMP4 treatment from days 26 to 35 maintained an ALDH⁺TH⁺ A9 population (Fig. 3c). This treatment did not efficiently induce ALDH activity or ALDH1A1 expression in A10-patterned cells, indicating a subtype-specific effect on A9 identity maintenance (Fig. 3d and Extended Figs. 5b,c).

We next examined ESRRB signaling, as ESRRB has been linked to mitochondrial oxidative phosphorylation programs and emerged from our multiome analysis as a candidate positive regulator of A9 development^46^. Addition of DY131 (2 μM), an ESRRB agonist, to Activin A and BMP4 from days 26 to 35 further increased ALDH expression by approximately 7% (Figs. 3e,f and Extended Fig. 5d).

We then validated this three-factor A9 maintenance condition in purified day 25 ALDH⁺ A9 cells. As negative controls, we included LDN-patterned A10-enriched cells and ALDH⁻ cells isolated from A9-enriched cultures (Fig. 3g). Following treatment with Activin A, BMP4 and DY131, ALDH activity was maintained in approximately 65–80% of A9 DDC⁺/TH⁺ neurons at day 35 and later stages (Figs. 3h,i and Extended Fig. 5e–g). By contrast, ALDH1A1 was not significantly induced in control populations or maintained in A9 cells cultured without the maintenance condition (Fig. 3i and Extended Fig. 5f–j). Consistent with preserved A9 identity, CALB1 expression remained significantly lower under the A9-maintenance condition than in A10 cultures (Fig. 3i). These findings were reproduced in an independent iPSC line, MSK-SRF001, in addition to KOLF2.1J and H9 lines (Extended Fig. 5k,l). Together, these results establish a postmitotic maintenance strategy that stabilizes ALDH1A1-expressing A9 identity after subtype specification, overcoming the loss of a key *in vivo* A9 feature during prolonged *in vitro* maturation.

### In vitro-generated mDA neuron subtypes are transcriptionally distinct

We next performed multiplexed scRNA-seq on day 36 populations, including purified A9 DA neurons maintained with Activin A and BMP4 (A+_AB4) or with additional DY131 (A+_AB4&DY), purified A10 DA neurons (LDN), LDN cells treated with DY131 (LDN_DY), and an intermediate population derived from day 25 ALDH⁻ cells isolated from A9-enriched cultures (Fig. 4a). hPSC-derived DA populations expressed canonical mDA markers, including *LMX1A*, *LMX1B* and *FOXA1*. We first classified cells as *ALDH1A1*⁺ A9 or *CALB1*⁺ A10 subtypes, and then assigned maturation states: mature DA neurons expressed high *TH* and *NR4A2*, progenitors showed low *TH/NR4A2* but high *SOX2* and *SOX9*, and intermediate clusters were annotated as early DA neurons (Figs. 4a,b and Extended Fig. 6a). UMAP analysis indicated that DY131 treatment did not alter DA subtype composition (Extended Fig. 6b).

**Fig. 4.**
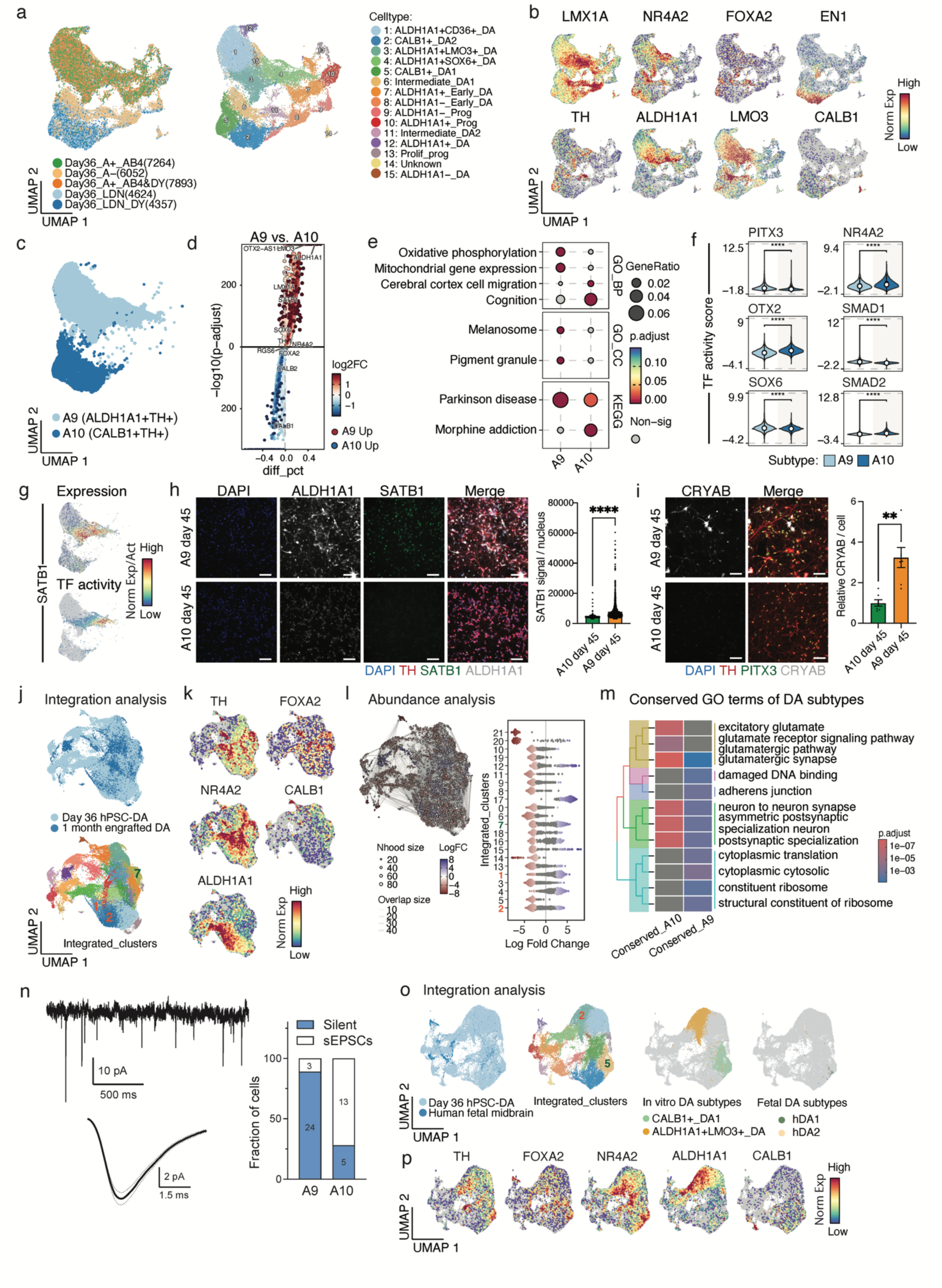
In vitro-generated mDA neuron subtypes are transcriptionally distinct. **a**, UMAP of scRNA-seq profiles from D36 hPSC-derived mDA populations colored by differentiation condition (left) and annotated cell type (right). **b,** UMAP feature plots showing expression of the indicated genes. **c,** UMAP highlighting grouped ALDH1A1+TH+ (A9) and CALB1+TH+ (A10) dopaminergic clusters. **d,** Volcano plot of differential gene expression between pseudo-bulk A9 and A10 groups. **e,** Pathway enrichment analysis of A9- and A10-enriched gene sets. **f,** Transcription factor (TF) activity scores for the indicated factors across clusters. **g,** SATB1 expression (top) and inferred TF activity (bottom) across the indicated populations. **h,** Representative IF images of SATB1 and ALDH1A1 in D45 A9 and A10 cultures (left), with quantification of nuclear SATB1 signal intensity (right). Student’s *t*-test; *n* = 3; scale bars, 100 µm; error bars, s.e.m. **i,** Representative IF images of CRYAB, TH and PITX3 in D45 A9 and A10 cultures (left), with quantification of relative CRYAB expression (right). Student’s *t*-test; *n* =5; scale bars, 100 µm; error bars, s.e.m. **j,** UMAP of CCA-integrated scRNA-seq data from D36 hPSC-mDA subtypes and 1-month grafted hPSC-mDA populations. **k,** UMAP feature plots showing expression of the indicated genes in the integrated dataset. **l,** Abundance analysis of integrated datasets across the indicated cell types/states. **m,** Pathway enrichment analysis of conserved A9 and A10 gene sets shared between D36 in vitro data and 1-month graft data. **n,** Whole-cell patch-clamp recordings showing silent neurons and those receiving spontaneous excitatory postsynaptic currents (sEPSCs), with associated quantification. **o,p,** UMAP of CCA-integrated D36 hPSC-mDA subtypes with human fetal mDA subtypes (o), and expression of the indicated mDA genes in the integrated dataset (p).

We first compared aggregated *TH⁺ALDH1A1⁺*(A9) and *TH⁺CALB1⁺* (A10) mDA neurons, which revealed distinct transcriptional signatures (Fig. 4c). Differential expression analysis (DEG) showed higher expression of *ALDH1A1, LMO3,* and *SOX6* in the *TH⁺ALDH1A1⁺* population, and higher expression of *CALB1* and *CALB2* in the *TH⁺CALB1⁺* population (Fig. 4d and Table S5). Gene ontology analysis further revealed subtype-relevant functional programs: A9 neurons were enriched for oxidative phosphorylation and pigment granule formation, hallmarks of SNpc DA neurons, whereas A10 neurons showed enrichment for cognition-and addiction-related processes, consistent with mesocorticolimbic function (Fig. 4e)^1^. A9 neurons also express higher levels of L-type Ca^2+^ channels genes *CACNA1D* and *CACNA2D1*, which are linked to the pacemaking properties of this subtype (Extended Fig. 6c)^47,48^.

Transcription factor (TF) activity analysis using decoupleR further uncovered subtype-specific regulatory programs consistent with prior murine studies: *SOX6, PITX3*, and *SMAD1* showed higher predicted activity in hPSC-derived A9 neurons, whereas *OTX2* and *SMAD2* activities were enriched in A10 neurons (Fig. 4f and Table S6)^49,50^. We next validated selected A9-enriched markers. SATB1, a regulator implicated in mDA neuron senescence and aging ^51,52^, showed higher expression and predicted TF activity in A9 neurons (Figs. 4g,h and Extended Fig. 6d). CRYAB, which is elevated in the substantia nigra of PD patients^53^, was likewise enriched in A9 neurons (Fig. 4i).

Finally, we benchmarked our hPSC-derived DA subtypes against published *in vitro* DA neuron datasets. CCA integration with iPSC- and hPSC-derived DA populations from Nishimura et al. showed that *CALB1* and *LMO3* were detectable across datasets, whereas robust *ALDH1A1* expression was largely restricted to the A9 populations generated using our method (Extended Figs. 6e-g)^54,55^. Comparison with assembloid-derived DA neurons from Reumann et al. similarly showed that our *ALDH1A1***⁺** populations expressed higher levels of A9 markers, including *ALDH1A1* and *LMO3* (Extended Fig. 6h–j)^56^.

Together, these analyses show that our strategy yields transcriptionally distinct human A9 and A10 mDA neurons with subtype-appropriate functional and regulatory features, and produces a more robust *ALDH1A1*⁺ A9 state than existing *in vitro* DA neuron systems.

### Transcriptomic similarity between in vitro-generated mDA neuron subtypes and in vivo-identified counterparts

To further assess the transcriptional fidelity of hPSC-derived DA subtypes and their similarity to *in vivo* counterparts, we performed CCA integration analyses comparing our *in vitro* hPSC-derived DA neurons with (1) hPSC-derived DA neurons grafted into the mouse striatum and (2) human fetal DA subtypes^2,8,55^. Integrated analysis of the *in vitro* and engraftment datasets showed that *CALB1*⁺ hPSC-derived DA neurons co-localized with graft-derived A10 neurons in cluster 7, whereas *ALDH1A1⁺LMO3⁺*hPSC-derived DA neurons clustered with graft-derived A9-like neurons in clusters 1 and 2 (Figs. 4j,k and Extended Figs. 7a,b). These findings were supported by MiloR-based relative abundance analysis^57^, which indicated that these clusters comprised both *in vitro*- and graft-derived cells, consistent with strong transcriptional similarity between hPSC-derived and *in vivo* DA subtypes (Fig. 4l).

To identify conserved molecular features of A9- and A10-like neurons, we performed DEG analysis on cells from clusters 1, 2, and 7, focusing on genes upregulated in either subtype irrespective of origin (*in vitro* vs grafted). GO enrichment analysis of these conserved DEGs revealed that A9-like neurons were enriched for ribosomal gene programs, whereas A10-like neurons upregulated genes associated with excitatory synapses and glutamate receptor signaling (Fig. 4m and Table S7)^58^. This aligns with the reported co-release of dopamine and glutamate by VTA DA neurons^59,60^. Consistent with this, patch-clamp recordings revealed more frequent spontaneous excitatory postsynaptic currents (sEPSCs) in A10 cultures, indicating stronger glutamatergic input (Fig. 4n).

Finally, integration of our hPSC-derived DA neurons with fetal midbrain datasets revealed strong transcriptional similarity between *in vitro*-generated *ALDH1A1⁺LMO3⁺* neurons and fetal hDA2 (*ALDH1A1⁺)* populations, as well as between *CALB1⁺* neurons and fetal hDA1 (*CALB1⁺)* populations (Figs. 4o,p and Extended Figs. 7c,d). Together, these results support the generation of authentic, human fetal-like A9 and A10 DA neuron subtypes from hPSCs.

### CREB signaling is required for the differentiation of hPSC-derived A10 mDA neurons

Leveraging these computational analyses, we tested a subtype-specific role for CREB signaling in A10 development. Multiome profiling nominated *CREB5* as a candidate positive regulator of A10 differentiation. Using hdWGCNA, we identified subtype-enriched gene modules in day 36 mDA neurons (Extended Figs. 7e,f)^61^, including an A9-enriched module 5 that overlapped with genes expressed in the human fetal hDA2 *ALDH1A1*⁺*LMO3*⁺ cluster (Extended Figs. 7g,h and Table S8), and an A10-enriched module 7 (Extended Figs. 7e–i). GO analysis revealed opposing CREB pathway signatures: module 7 was enriched for positive regulation of CREB signaling, whereas module 5 was enriched for negative regulation (Extended Figs. 7i–k). Consistent with these findings, *in silico* knockout of *CREB5* preferentially dysregulated genes expressed in *CALB1*⁺*TH*⁺ neurons and disrupted the A10 gene regulatory network, whereas knockout of *ID4*, a predicted A9 regulator, primarily affected the *ALDH1A1*⁺*TH*⁺ population (Extended Fig. 7l)^62^. Functionally, CREB inhibition (1 μM 666-15) reduced PITX3 expression in A10-patterned cultures at day 25 and decreased TH expression by day 30, with no comparable effect in A9 cultures (Extended Figs. 7m–o). These results identify a selective requirement for CREB signaling in human A10 mDA neuron development.

### Biochemical and electrophysiological distinctions between hPSC-derived mDA neuron subtypes

Guided by GSEA showing enrichment of pigment biogenesis and mitochondrial pathways in *ALDH1A1*⁺ A9 neurons, with relative downregulation in *CALB1*⁺ A10 neurons (Fig. 5a), we assessed functional differences between these hPSC-derived DA subtypes at days 55–60. Electron microscopy revealed more elaborated mitochondrial cristae in A9 neurons, whereas A10 cells contained more enlarged double-membrane organelles, indicating less functional mitochondria (Fig. 5b). Consistent with these findings, Seahorse analysis revealed substantially higher basal and maximal mitochondrial respiration in A9 neurons than in A10 neurons (basal respiration, 177 versus 3.5; maximal respiration, 248 versus 10)^63^, indicating greater oxidative metabolic capacity in A9 (Fig. 5c).

**Fig. 5.**
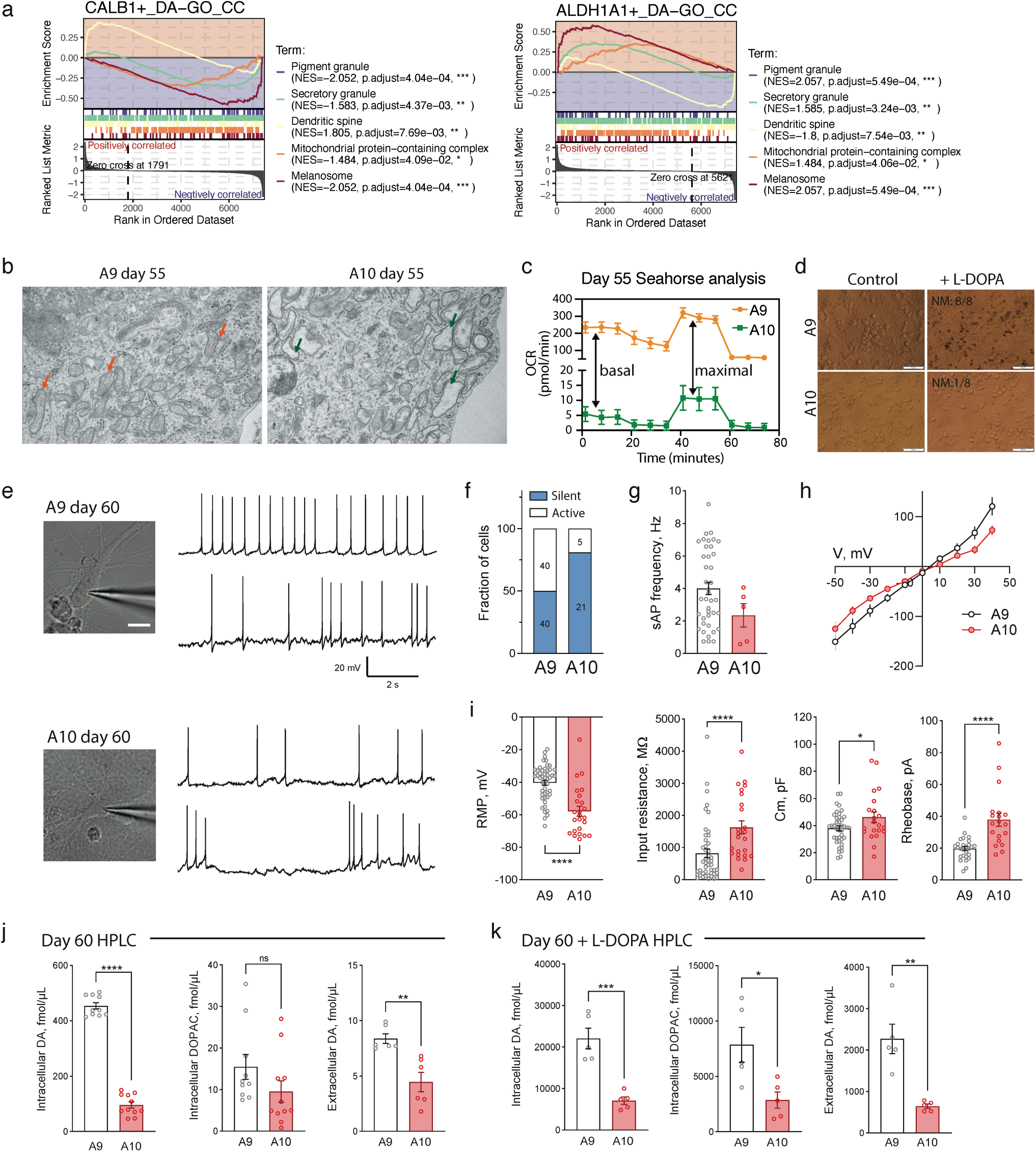
Biochemical and electrophysiological distinctions between hPSC-derived mDA neuron subtypes. **a**, GSEA showing enrichment and depletion of selected pathways in CALB1+ DA and ALDH1A1+ DA clusters. **b,** Representative electron microscopy images of D55 A9 and A10 neurons, with mitochondria highlighted. Scale bar, 600 nm. **c,** Seahorse extracellular flux analysis of D55 A9 and A10 mDA neurons. **d,** Representative images of A9 and A10 mDA neurons treated with L-DOPA. Scale bars, 50 µm. **e,** Brightfield images of patched A9 and A10 neurons (left; Scale bar, 10 µm) and representative whole-cell recordings of spontaneous activity (right). **f,** Fraction of spontaneously active versus silent neurons; numbers of recorded neurons are indicated within bars. Chi-square test. **g,** Action potential firing frequency in spontaneously active neurons. Mann–Whitney test; not significant. **h,** Current–voltage relationship for A9 and A10 neurons recorded in voltage-clamp mode. Two-way ANOVA; not significant. **i,** Resting membrane potential, input resistance, membrane capacitance and rheobase for A9 and A10 neurons. Mann–Whitney test. **j,k,** HPLC quantification of intracellular dopamine (DA), intracellular DOPAC and extracellular DA in the absence (j) or presence (k) of L-DOPA treatment. Mann–Whitney test. All error bars in the figure represent s.e.m.

We next assessed pigmentation capacity following L-DOPA treatment (50 µM, 4–7 days)^64^. Pigment accumulation occurred predominantly in A9 neurons (8/8 A9 cultures versus 1/8 A10 cultures), indicating subtype-biased neuromelanin-like pigment formation *in vitro* (Fig. 5d). This pigmentation was accompanied by neuronal degeneration in A9 cultures (Extended Fig. 8a), potentially reflecting the intrinsic vulnerability of this subtype to dopamine-mediated oxidative stress.

Patch-clamp recordings showed that A9 neurons were more electrophysiologically active than A10 neurons, firing at approximately 4 Hz versus 2 Hz (Figs. 5e–g), consistent with the autonomous low-frequency pacemaking activity characteristic of SNpc neurons^65^. A9 neurons also exhibited a higher resting membrane potential and lower input resistance, membrane capacitance and rheobase than A10 neurons (Figs. 5h,i). The reduced rheobase is consistent with greater intrinsic excitability, in line with prior evidence that SNc- and VTA-associated dopaminergic populations differ in their passive and active membrane properties^66,67^. High-Performance Liquid Chromatography (HPLC) analysis further showed higher intracellular and extracellular dopamine levels in A9 neurons under basal conditions and after L-DOPA treatment (Figs. 5j,k). Together, these findings show that hPSC-derived A9 and A10 neurons acquire distinct subtype-relevant functional properties, with A9 neurons recapitulating key features of SNpc DA neurons.

### Subtype-specific neuromelanin and IFN-γ converge to activate microglial inflammation and MHC-I antigen presentation in A9 mDA neurons

Inflammatory interactions with microglia and T cells are thought to contribute to the selective vulnerability of SNpc dopaminergic neurons^68–74^. Studies have shown that microglia can be activated by neuromelanin released from A9 neurons^75,76^, leading to secretion of pro-inflammatory factors, and that CD8⁺ T cells can recognize MHC-I–presented antigens on A9 neurons and target them^68,70,77,78^. To model these interactions, we treated hPSC-derived microglia with neuromelanin-like structures generated from A9 cultures (Figs. 6a,b)^79^. This induced secretion of pro-inflammatory cytokines, including IL-1β, TNF-α, IL-8 and IL-6 (Fig. 6c).

**Fig. 6.**
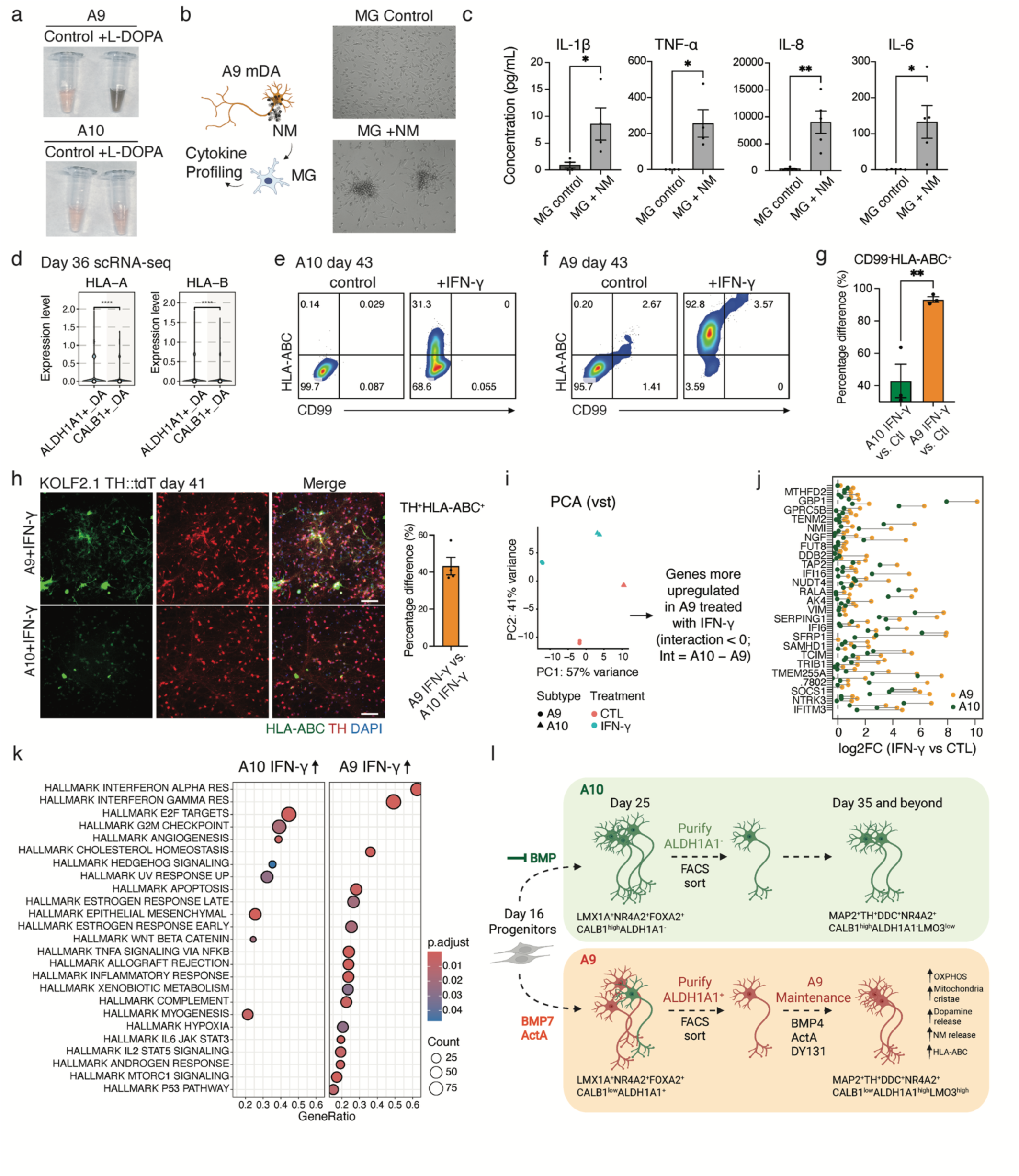
Subtype-specific neuromelanin and IFN-γ converge to activate microglial inflammation and MHC-I antigen presentation in A9 mDA neurons. **a**, Representative media color change following L-DOPA treatment in A9 cultures. **b,** Schematic of treating hPSC-derived microglia with neuromelanin (NM)-like structures produced by hPSC-derived A9 neurons. **c,** Secretion of the indicated proinflammatory cytokines by hPSC-microglia following NM-like structure exposure. Student’s *t*-test; *n* = 4–5 independent differentiations; error bars, s.e.m. **d,** Expression of *HLA-A* and *HLA-B* in D36 ALDH1A1+ DA and CALB1+ DA clusters. Wilcoxon test. **e,f,** Flow cytometric of CD99 and HLA-ABC expression in hPSC-A10 (e) and hPSC-A9 (f) cultures ± IFN-γ. **g,** Quantification of CD99−HLA-ABC+ cells with and without IFN-γ across subtypes. Student’s *t*-test; *n* = 3 differentiations; error bars, s.e.m. **h,** Representative IF images of HLA-ABC and TH in the indicated populations (left), with quantification of TH+HLA-ABC+ cells in IFN-γ–treated A9 versus A10 cultures (right). Scale bars, 100 µm. **i,** PCA of bulk RNA-seq profiles from A9 and A10 cultures ± IFN-γ. **j,k,** Genes preferentially upregulated by IFN-γ in A9 relative to A10 (j), and pathway enrichment of IFN-γ–responsive gene sets in each subtype (IFN-γ versus control) (k). **l,** Working model summarizing differentiation of A9 and A10 subtypes from hPSCs.

We next assessed HLA-ABC/MHC-I expression in hPSC-derived mDA subtypes. Although baseline expression was low, day 36 scRNA-seq revealed higher HLA-A and HLA-B expression in *ALDH1A1*⁺ mDA neurons (Fig. 6d). Notably, FACS showed that IFN-γ treatment (20 ng/mL) robustly upregulated HLA-ABC in CD99⁻TH⁺ A9 neurons, with a significantly milder effect in A10 neurons (Figs. 6e-g and Extended Fig. 8b); IF analysis showed a similar trend (Fig. 6h). In contrast, MHC class II expression was not induced in mDA neurons following IFN-γ treatment (Extended Data Fig. 8c).

To define this subtype-selective inflammatory response, we performed bulk RNA-seq on day 42 A9 and A10 neurons with or without IFN-γ treatment (Fig. 6i and Extended Fig. 8d). GSEA using an IFN-γ-responsive gene set from Hobson et al. confirmed pathway activation in both subtypes (Extended Fig. 8e)^71^. However, A9 neurons mounted a stronger antigen-presentation response, with greater induction of MHC-I genes (*HLA-B*, *HLA-C*), immunoproteasome components (*PSMB* genes) and antigen-processing machinery (*TAP1*, *TAP2*, *ERAP1*, *ERAP2*) (Extended Fig. 8f)^80^. Genes preferentially induced in A9 relative to A10 neurons were enriched for interferon-α/γ response, inflammation, apoptosis and p53-associated pathways (Figs. 6j,k). A9 neurons also more strongly upregulated T cell-recruiting chemokines (*CCL25, CXCL10, MIF*) and the microglia-recruiting chemokine *CX3CL1* (Extended Fig. 8g)^68^.

IFN-γ treatment was additionally associated with reduced expression of core dopaminergic markers (*NR4A2, DDC, FOXA2, TH*) and *CALB1*. Notably, IFN-γ differentially affected A9-associated genes across subtypes: *ALDH1A1* and *RGS6* were downregulated in A9 neurons but upregulated in A10 neurons (Extended Fig. 8h).

Together, these findings establish a human subtype-resolved neuroimmune platform linking intrinsic A9 identity to microglial activation and heightened antigen-presentation responses, potentially contributing to its selective vulnerability in PD.

## Discussion

We have established a stepwise framework for generating transcriptionally, molecularly, and functionally distinct A9- and A10-like mDA neurons from hPSCs (Fig. 6l). Although floor plate-based protocols have markedly improved the quality, graftability, and translational potential of hPSC-derived mDA neurons, they have largely focused on producing pan-mDA populations^6,7,81–83^. The controlled derivation of developmentally distinct A9- and A10-like subtypes has remained a major unmet goal. Our work addresses this gap by showing that human mDA subtype identity can be prospectively specified at the progenitor stage and reinforced during neuronal maturation, yielding cells that capture key *in vivo* subtype-associated transcriptional, metabolic, functional, and immune-responsive features. This framework moves the field beyond generic dopaminergic differentiation toward subtype-resolved human systems for modeling selective vulnerability in diseases, neural circuitry, and precision cell therapy in PD.

A central advance is the demonstration that mDA subtype generation involves two separable phases: early subtype specification and later subtype maintenance. We show that Activin A and BMP7 act during the progenitor-to-neuron transition to bias cells toward an A9-like trajectory, whereas maintenance of postmitotic A9 features requires a distinct regime involving ESRRB activation together with Activin A and BMP4, rather than BMP7. This stage-specific logic suggests that early patterning alone is insufficient to preserve a broader A9-like program through maturation, and may help explain why prior protocols cannot robustly stabilize subtype identity^21^.

Our findings identify BMP signaling as a determinant of A9 versus A10 divergence and extend emerging evidence that TGF-β superfamily pathways shape multiple stages of dopaminergic development. *In vivo* studies have implicated BMP5/6/7 and SMAD1 activity in ventral midbrain dopaminergic neurogenesis and substantia nigra neuron development^84,85^. More recently, Terauchi *et al.* showed that BMP6/BMP2–SMAD1 and TGF-β2–SMAD2 signaling differentially support nigrostriatal and mesolimbic dopaminergic synapse formation, linking distinct SMAD axes to projection-specific circuit assembly^50^. Our work extends this concept to human subtype specification by showing that BMP activity is not merely permissive for mDA differentiation, but instructive for A9 fate bias, promoting ALDH1A1-associated A9 features while restraining CALB1-associated A10 identity. The distinct use of BMP7 during subtype induction and BMP4 during A9 maintenance further points to ligand- and stage-specific BMP logic. BMP7 may participate in an early ventral midbrain specification program, whereas BMP4 more effectively preserves postmitotic A9 features. Together, these findings suggest that TGF-β superfamily signals encode dopaminergic subtype properties through ligand identity, developmental timing, and cellular competence.

A second advance is the establishment of reporter-free strategies to isolate and track emerging human mDA subtypes. By combining CD56/CD171-based neuronal enrichment, CD99-based exclusion of less differentiated or TTR^+^ contaminating populations, and ALDH enzymatic activity, our approach physically resolves subtype-enriched cells without engineered reporters. This provides prospective access to A9- and A10-biased developmental intermediates and enables human subtype decisions to be studied dynamically rather than inferred from endpoint cultures.

The ability to generate and isolate subtype-enriched postmitotic A9- and A10-like neurons also has implications for cell replacement therapy. Current clinical and preclinical efforts have established the feasibility of transplanting hPSC- or iPSC-derived ventral midbrain dopaminergic progenitors, while recent work with hPSC-derived A10 neurons highlights the therapeutic relevance of subtype-specific dopaminergic populations^5,7,83^. Yet how graft subtype composition influences engraftment fidelity, circuit reconstruction, and functional outcome remains unclear. Our finding that purified postmitotic A9- and A10-like neurons survive after striatal transplantation and retain subtype-associated features *in vivo*, including differential GIRK2 and CALB1 expression, provides a crucial framework for addressing this question. Although functional benefit was not assessed here, this platform enables future testing of whether A9-enriched grafts more faithfully restore the dopaminergic populations preferentially lost in PD.

Our system also enables subtype-resolved modeling of neuroinflammatory interactions relevant to PD. We show that hPSC-derived A9 neurons form neuromelanin-like structures that trigger pro-inflammatory cytokine secretion from hPSC-derived microglia, extending prior observations from human tissue and model systems into a defined human cellular platform. Recent work has established the feasibility of modeling T cell–midbrain interactions using human midbrain organoids, showing that activated peripheral T cells infiltrate organoid tissue and promote neuronal loss; however, this system used polyclonally activated T cells, lacked detectable MHC-I expression in the organoids, and therefore primarily modeled MHC-independent cytotoxicity^86^. By contrast, we show that hPSC-derived A9-like neurons exhibit stronger IFN-γ-induced upregulation of MHC-I and antigen-processing machinery than A10-like neurons, revealing a previously unrecognized subtype-selective immune responsiveness in human mDA neurons. These findings link A9 identity to neuronal states that may facilitate antigen-dependent T cell recognition and provide a more mechanistically defined, PD-relevant framework for modeling immune-mediated selective vulnerability.

Defined A9- and A10-like neurons should further improve the biological specificity of human circuit and assembloid models. Existing stem cell-based dopaminergic circuits often use poorly resolved mDA populations despite the distinct nigrostriatal, mesolimbic, and mesocortical pathways formed by A9 and A10 neurons *in vivo*^56^. Our differentiation framework should enable more faithful pathway-specific models, pairing A9-like neurons with dorsal striatal targets and A10-like neurons with limbic or cortical targets, and may help distinguish mechanisms underlying motor versus non-motor dopaminergic dysfunction.

The current system does not fully capture all features of mature *in vivo* mDA subtypes, and the identification of an *ALDH1A1*-negative, pan-mDA-positive intermediate population within A9-enriched cultures suggests that human subtype maturation may proceed through a more complex continuum than a simple binary A9/A10 branch. These cells may represent a delayed maturation state, an incompletely resolved subtype intermediate, or a parallel mDA-like identity that warrants further study through lineage tracing and integration with developing human midbrain and annotated *in vitro* references^87,88^.

Together, our findings establish a mechanistically informed platform for specifying, maintaining, isolating, and interrogating human mDA neuron subtypes. By resolving subtype identity across developmental, transplant, and immune dimensions, this work provides a foundation for precision models of the human SNpc and VTA, with implications for PD modeling, circuit biology, and subtype-informed cell therapy.

## Methods

### hPSC culture

Human pluripotent stem cell lines (WA09 (H9; 46,XX), H9 PITX3::GFP, J1 (MRC5), MSK-SRF001, and KOLF2.1 TH::tdTomato) were maintained on Vitronectin (VTN-N; Thermo Fisher) coated plates in Essential 8 medium (Life Technologies, A1517001). Cells were dissociated with EDTA and passaged 1to 2 times before differentiation. All lines were cultured at 37 °C and 5% CO₂ and routinely tested for mycoplasma. All stem cell work was performed under protocols approved by the Tri-Institutional Stem Cell Initiative Embryonic Stem Cell Research Oversight Committee (Tri-SCI ESCRO).

### Directed differentiation into midbrain dopaminergic neuron subtypes from hPSCs

hPSCs were dissociated to single cells with Accutase and plated at 5 × 10^5 cells/cm^2 on Geltrex (Life Technologies, A1413201)–coated plates in Neurobasal medium supplemented with N2 (Stemcell Technologies), B27 (Life Technologies) and 2 mM L-glutamine, together with SHH C25II (500 ng/ml; R&D Systems), LDN193189 (250 nM; Stemgent), SB431542 (10 µM; R&D Systems), CHIR99021 (1 µM; R&D Systems) and ROCK inhibitor Y-27632 (10 µM; R&D Systems). This was designated day 0 of differentiation. ROCK inhibitor was removed from day 1, and cultures were maintained through day 3. From day 4 to day 7, CHIR99021 was increased to 6 µM . On day 7, SHH, LDN193189 and SB431542 were withdrawn (only 6 µM CHIR). On day 10, media was switched to Neurobasal/N2/B27/L-glutamine supplemented with BDNF (20 ng/ml; R&D Systems), ascorbic acid (0.2 mM; Sigma), GDNF (20 ng/ml; PeproTech), TGFβ3 (1 ng/ml; R&D Systems), dibutyryl cAMP (0.2 mM; Sigma) and CHIR99021 (3 µM). On day 11, cells were dissociated with Accutase and replated at high density (8.5 × 10^5 cells/cm^2) onto polyornithine (15 µg/ml; Sigma Aldrich)/laminin (1 µg/ml; R&D System)/fibronectin (2 µg/ml; Thermo Fisher)–coated plates in day 10 mDA differentiation medium (Neurobasal/N2/B27/L-glutamine with BDNF, ascorbic acid, GDNF, dibutyryl cAMP, CHIR99021 and TGFβ3). From days 12 to 16, cultures were maintained in Neurobasal/B27/L-glutamine supplemented with BDNF, ascorbic acid, GDNF, dibutyryl cAMP, TGFβ3, IWP2 (1 µM; Selleckchem) and FGF18 (100 ng/ml; PeproTech). On day 16, cells were dissociated and replated as in day 11, and maintained until day 25 in mDA differentiation medium supplemented with DAPT (10 µM; R&D Systems) (Neurobasal/B27/L-glutamine with BDNF, ascorbic acid, GDNF, dibutyryl cAMP, TGFβ3 and DAPT)^8^. For cryopreservation of mDA neuron precursors, day 16 cells were dissociated with Accutase to single cells, resuspended at 1.5 × 10^7 cells/ ml in STEM-CELLBANKER, and frozen using a controlled-rate freezer (Thermo Fisher). Subtype patterning was initiated on day 17: Activin A (10 ng/ml; R&D Systems) plus BMP7 (20 ng/ml; R&D Systems) was used to bias progenitors toward the A9 lineage, whereas LDN193189 (250 nM) was used to bias toward the A10 lineage. Media was changed every other day from days 17-25. On day 25, cells were dissociated with Accutase and replated at low density (2.5 × 10^5 cells/cm^2) in day 16 mDA differentiation medium and maintained until downstream assays. For pathway perturbation experiments, cultures were treated from day 17 to day 25 with Wnt C-59 (Cayman; 1 µM), the CREB inhibitor 666-15 (MCE; 1 µM), or the MEK inhibitor PD0325901 (Selleckchem; 10 µM), with fresh compounds added every other day. For A9 maintenance, beginning on day 26 (1 day after replating), cultures were supplemented with Activin A (10 ng/ml), BMP4 (R&D Systems; 20 ng/ml), and DY131 (MCE; 2 µM). To induce pigmentation in A9 cultures, levodopa (Selleckchem; 50 µM) was applied for 4–7 days, until the culture medium became dark brown. Details of all reagents are provided in Table S3.

### Generation and verification of the H9 PITX3::GFP reporter line

The H9 PITX3::GFP reporter line was generated using CRISPR/Cas9-mediated homology-directed repair (HDR)^89,90^. Briefly, a single-guide RNA (sgRNA) was designed to target a sequence proximal to the stop codon of the PITX3 gene and cloned into the pX330-U6-Chimeric_BB-CBh-hSpCas9 vector (Addgene plasmid, Cat#42230) to generate the sgRNA-expressing construct. A donor plasmid was constructed containing 550 bp left and right homology arms, with a P2A-H2B-GFP cassette and a floxed puromycin resistance cassette positioned between the homology arms. The sgRNA/Cas9 plasmid and donor plasmid were co-electroporated into H9 human embryonic stem cells using a Lonza 4D-Nucleofector with Solution Primary Cell P3 and pulse code CB-150. Puromycin selection was initiated 3 days post-electroporation and maintained for 4 days to enrich for donor knock-in cells. Single-cell clones were subsequently derived, and correct knock-in clones were identified by PCR screening and Sanger sequencing. Expression of the reporter gene was further validated using CRISPRa-mediated gene activation, as previously described. Briefly, the SAM-TET1 system together with an sgRNA targeting PITX3 (spacer sequence: CCAGACCCCTCCCTCAAAGG) was electroporated into PCR-positive clones to transiently activate endogenous PITX3 expression. GFP expression was detected 48 h post-electroporation, confirming functional reporter activation.

### RNA extraction and Real-time qRT-PCR

Total RNA was extracted using the RNAqueous-Micro Kit (Invitrogen) and treated with RNase-free DNase (Invitrogen). DNase-treated RNA was reverse transcribed using a mixture of oligo(dT) primers and random hexamers with iScript reverse transcriptase (Bio-Rad). Real-time PCR was performed using SsoFAST EvaGreen Mix (Bio-Rad) in a Bio-Rad CFX96 Thermal Cycler. Absolute copy numbers were calculated from standard curves generated with human genomic DNA, and relative expression was obtained by normalizing each target to TBP. Primer sequences are listed in Table S1.

### Immunohistochemistry

Cells were fixed in 4% paraformaldehyde (PFA; Affymetrix) in DPBS for 13 min at room temperature and washed with DPBS. Samples were permeabilized with 0.5% Triton X-100 in DPBS for 25 min and blocked for 1 hour in blocking buffer (2% BSA and 0.25% Triton X-100 in DPBS). Primary antibodies were incubated overnight at 4 °C in blocking buffer. After DPBS washes, samples were incubated for 1 hour at room temperature with Alexa Fluor–conjugated secondary antibodies (488, 555 or 647; Invitrogen) in blocking buffer. Samples were washed with DPBS and counterstained with DAPI (Sigma) for 5 min. Images were acquired on Olympus or Zeiss inverted fluorescence microscopes. For HLA-ABC and MHC-II immunostaining, cells were fixed with 4% paraformaldehyde without permeabilization. During blocking and primary antibody incubation, 0.1% Tween 20 was used in place of 0.5% Triton X-100. Antibody details and dilutions are provided in Table S2.

### Cell surface marker staining and flow cytometry analysis

Cells were dissociated to single cells with Accutase for 30 min at 37 °C and washed with DMEM base medium (Thermo Fisher). Cells were pelleted at 500g for 3 min, washed, resuspended, and stained with fluorophore-conjugated antibodies diluted in DMEM base medium for 20 min at room temperature. Stained cells were washed twice with DMEM, filtered through cell-strainer–cap tubes (Falcon), and analyzed on an LSRFortessa (BD Biosciences) flow cytometer. Data were processed in FlowJo. Unstained and secondary-only controls were included for gating. Antibody details and dilutions are provided in Table S2.

### Aldehyde dehydrogenase assay

The ALDEFLUOR™ assay (STEMCELL Technologies, 01700) and AldeRed® assay (Sigma-Aldrich, SCR150) were used to measure aldehyde dehydrogenase (ALDH) activity in A9-patterned and sorted cultures^31^. Cells were dissociated as described in the “Flow Cytometry” section and incubated in ALDEFLUOR assay buffer containing ALDEFLUOR substrate (1:1000), or in AldeRed buffer containing verapamil and AldeRed substrate (1:1000), for 30–40 min at 37 °C in the dark. As a negative control, an aliquot of substrate-treated cells was incubated with the ALDH inhibitor DEAB (1 nM). Reactions were stopped by washing cells with cold wash medium (Neurobasal supplemented with 10% ALDEFLUOR/AldeRed assay buffer), and cells were maintained in this cold wash medium during analysis. For sorting ALDH⁺ A9 populations, stained cells were kept in cold Neurobasal medium containing 10% ALDEFLUOR/AldeRed assay buffer throughout sorting. Kit information is listed in Table S4.

### Microglia differentiation and inflammatory cytokine analysis

Microglia were generated based on a previously published protocol^79^. Briefly, wildtype H9 (WA09) hES cells were dissociated into a single-cell suspension on Day 0 using 0.01% Trypsin/EDTA. 40,000 cells per cm2 were seeded onto Matrigel-coated plates in Essential 8 (E8) medium with Activin A (7.5 ng/ml; R&D), BMP4 (30 ng/ml; R&D), CHIR 99021 (3 μM; Tocris Bioscience), and Y-27632 (10 μM, Invitrogen). After 18 hours, medium was changed to Essential 6 (E6) with Activin A (10 ng/ml), BMP4 (40 ng/ml), and IWP2 (2 μM; Selleckchem). On day 2, E6 medium was changed with Activin A (10 ng/ml), BMP4 (40 ng/ml), IWP2 (2 μM), and FGF2 (20 ng/ml; R&D). On day 3, cells were dissociated with Accutase and re-seeded at 60,000 cells per cm2 in Matrigel coated plates in E6 medium with Y-27632 (10 μM), VEGF (15 ng/ml; R&D) and FGF2 (5 ng/ml). On Day 4, medium was changed to Day 3 media without Y-27632. On day 5 and 6, E6 medium was changed with VEGF (15 ng/ml), FGF2 (5 ng/ml), SCF (200 ng/ml; R&D) and IL-6 (20 ng/ml; R&D), On day 7 and 9, E6 medium was changed with SCF (100 ng/ml), IL-6 (10 ng/ml), TPO (30 ng/ml; R&D), and IL-3 (30 ng/ml; R&D). On day 10, the round cells were harvested and directly plated into RPMI media (Gibco) containing 10% FBS (R&D), L-glut (Gibco), Pen/Strep (Gibco), IL-34 (100 ng/ml; R&D), and M-CSF (10 ng/ml; R&D) for 10 days until round cells were adherent. Matured microglia were replated at 40,000 cells per cm2 in 96-well plates and cultured for 48 hours, after which the cells were switched to RPMI medium without FBS for 2 days. For testing microglial inflammatory response towards neuromelanin, medium from a 96-well of A9 dopaminergic neurons (∼80,000 cells per well) treated with 50 μM levodopa (Selleckchem) for 4 to 7 days was collected and the sedimented neuromelanin was added to one 96-well of microglia. After 48 hours, the conditioned medium was collected and subjected to inflammatory cytokine analysis using the LEGENDPlex multi-analyte flow assay kit (BioLegend, 740808) according to the manufacturer’s instruction. Analysis was done at legendplex.qognit.com.

### Seahorse extracellular flux analysis

Dopaminergic neurons were plated on Seahorse XF96 microplates (Agilent, 103794-100) coated with poly-D-lysine, fibronectin and laminin at a density of 80K cells/well. The cells were then analyzed on days 55 of differentiation. Seahorse XF96 sensor cartridges were hydrated overnight at 37°C in a non-CO₂ incubator using XF calibrant solution (Agilent, 100840-000). On the day of assay, cells were equilibrated in DMEM-based Seahorse XF medium (Agilent,103680-100) supplemented with glucose (10 mM; Agilent, 103577-100), pyruvate (1 mM ; Agilent, 103578-100), and glutamine (2 mM; Agilent, 103579-100) (pH 7.4) for 45–60 min at 37°C. Oxygen consumption rate (OCR) was measured using the Seahorse XF Mito Stress Test (Agilent, 103015-100) with sequential injections of oligomycin (1 µM), FCCP (2 µM), and rotenone/antimycin A (0.5 µM each). Measurements were acquired using repeated mix–wait–measure cycles (3–3–3 min; three cycles per condition). OCR values were normalized to nuclei counts (redo-imaging).

### Electron microscopy

Cultures were fixed in 2% glutaraldehyde in 0.08 M sodium cacodylate buffer (pH 7.2) 2 mM CaCl_2_ for 4 hs at room temperature followed by overnight fixation at 4°C, postfixed in 1% osmium/0.8% potassium ferricyanide in 0.1 M cacodylate buffer, followed by post-staining in 1% uranyl acetate in 0.05 M maleate buffer pH 5.2, dehydration in an ethanol series, and embedding in Eponate 12 (Ted Pella, Inc). Ultrathin sections (60–65 nm) were stained with uranyl acetate and lead citrate, and images were acquired using a Tecnai G2-Spirit transmission electron microscope (FEI, Hillsboro, Oregon) operated at 120 kV, equipped with an AMT BioSprint29 digital camera (AMT, Danvers, MA).

### Sample preparation, single-cell library generation, hashtag demultiplexing and data processing

Cells were dissociated to single cells using Accutase as described above and stained with DAPI. DAPI-negative (live) cells were sorted on a FACSAria Fusion (BD Biosciences) at the MSKCC Flow Cytometry Core. To minimize batch effects, samples collected at the same differentiation stage (for example, all day 25 conditions) were processed in parallel: they were prepared, hashtag-labeled, FACS-purified, pooled, and sequenced together. Hashtag-labeled samples from each stage were pooled and sequenced on a single lane. Single-cell 3ʹ gene expression libraries were generated from the sorted suspensions using the 10x Genomics Chromium 3ʹ GEM-X kit according to the manufacturer’s instructions. For scRNA-seq, day 25 and day 36 samples were generated from the H9 PITX3::GFP and KOLF2.1 TH::tdTomato reporter lines, respectively. Individual conditions were labeled with TotalSeq™-A hashtag oligonucleotides (BioLegend, Series A) according to the manufacturer’s instructions, pooled at relatively equal proportions, and processed together for 10x Genomics single-cell library preparation and sequencing. Hashtag reads were used to demultiplex pooled libraries; cells classified as “unmapped” were excluded from downstream analyses. Hashtag assignments were as follows. Day 25 pool (H9 PITX3::GFP-derived): A0253 (TTCCGCCTCTCTTTG): Day25_ACTA_BMP7_MI, A0254 (AGTAAGTTCAGCGTA): Day25_LDN, A0259 (CAGTAGTCACGGTCA): Day25_LDN_MI, A0251 (GTCAACTCTTTAGCG): Day25_Control, A0252 (TGATGGCCTATTGGG): Day25_ ACTA_BMP7. Day 36 pool (KOLF2.1 TH::tdTomato): A0253 (TTCCGCCTCTCTTTG): Day36_LDN, A0254 (AGTAAGTTCAGCGTA): Day36_LDN_DY, A0259 (CAGTAGTCACGGTCA): Day36_int_AB4_DY, A0251 (GTCAACTCTTTAGCG): Day36_A+_AB4, A0252 (TGATGGCCTATTGGG): Day36_A−. Demultiplexing and alignment to the GRCh38 reference genome was performed with the CellRanger pipeline (v7.1.0). Data analysis was performed with R (v4.4.2) using Seurat_(v5.0.3).

Hashtag oligonucleotide (HTO) counts from TotalSeq sample-barcoding antibodies were used to demultiplex pooled 10x Genomics single-cell libraries. Antibody details are provided in Table S2. After alignment and quantification of gene expression, HTO feature-barcode count matrices were generated from the 10x Genomics Feature Barcode pipeline using the corresponding TotalSeq hashtag reference described above. The filtered gene expression matrix and the filtered HTO feature-barcode matrix were processed with Seurat. HTO assays were added to the Seurat object and normalized using centered log-ratio (CLR) normalization across cells (NormalizeData, assay = “HTO”, normalization.method = “CLR”). Demultiplexing was performed using Seurat’s HTODemux function with the CLR-normalized HTO signal (HTODemux, assay = “HTO”, positive.quantile = 0.95).

Raw 10x Genomics gene expression matrices were processed using Seurat. Low-quality cells were removed on a per-sample basis using standard QC metrics, including the number of detected genes per cell (nFeature_RNA), total UMI counts (nCount_RNA), and the fraction of mitochondrial reads (percent.mt). Specifically, cells were retained if their nFeature_RNA and percent.mt values fell within the 5^th^ to 95^th^ percentile of the corresponding sample distributions. Data were normalized and variance-stabilized using SCTransform (Seurat; SCTransform, assay = “RNA”) with default parameters unless noted. Putative doublets were identified using DoubletFinder (v2.0.6) on the normalized data.

For each sample, an expected doublet rate of 3% was assumed and the number of predicted doublets was set to nExp = round(0.03 × Ncells), where Ncells denotes the number of post-QC cells; predicted doublets were removed prior to downstream analyses. Following SCTransform, principal component analysis (PCA) was performed on Pearson residuals, and cells were embedded using UMAP (RunUMAP, dims = 1:35). Nearest-neighbor graphs were constructed (FindNeighbors, dims = 1:35) and clustering was performed using the Louvain algorithm (FindClusters, resolution = 0.6), following standard Seurat SCTransform workflows. Data visualization was performed with SCP (v0.5.6).

### Integration of multiple datasets

To compare our dataset with external in vitro and in vivo single-cell transcriptomic references, the corresponding Seurat objects were merged and integrated using Seurat’s anchor-based integration workflow, leveraging canonical correlation analysis (CCA) with SCTransform normalization^55^. Briefly, the two datasets were merged into a single object and then split by sample group (metadata field “Data Source”). Each subset was independently normalized and variance-stabilized using SCTransform (SCTransform, default parameters). A shared set of integration features (n = 2,000) was selected using SelectIntegrationFeatures, and SCTransform-based integration preparation was performed with PrepSCTIntegration. Integration anchors were identified across datasets using FindIntegrationAnchors with normalization.method = “SCT” and the selected integration features, and an integrated expression matrix was generated using IntegrateData (normalization.method = “SCT”). The integrated object was then subjected to principal component analysis (RunPCA) and the number of principal components used for downstream analyses was selected based on elbow plot inspection (typically the first 30 PCs). Cells were embedded in two dimensions using UMAP (RunUMAP, reduction = “pca”, dims = 1:30). A shared nearest-neighbor graph was constructed (FindNeighbors, dims = 1:30) and unsupervised clustering was performed (FindClusters, resolution = 0.6). For clustering and visualization on the integrated space, the default assay was set to the integrated assay; for downstream gene-level analyses, the default assay was set back to the SCTransform assay as indicated (“SCT” assay). Integrated embeddings were visualized by dataset/stage using UMAP projections.

### Transcription factor activity inference

Transcription factor (TF) activities were inferred from single-cell RNA-seq data using decoupleR ((v 2.9.7)^49^. A curated human TF–target regulatory network was obtained from the CollecTRI resource via decoupleR::get_collectri (organism = “human”, split_complexes = FALSE), which provides signed TF–target interactions (mode of regulation, mor). Gene expression values were extracted from the Seurat object as a gene-by-cell matrix using the normalized RNA data slot (RNA assay) and converted to a numeric matrix.

TF activities were estimated using the univariate linear model (ULM) implemented in decoupleR (run_ulm) with the CollecTRI network, specifying the regulator column as .source = “source”, the target column as .target = “target”, and the sign column as .mor = “mor”. Regulons with fewer than five target genes were excluded (minsize = 5). The resulting TF activity scores were reshaped to a TF-by-cell matrix and stored as a new assay in the Seurat object by creating a Seurat assay (CreateAssayObject) and adding it to the object (assay name: “tfsulm”). Downstream visualization and differential TF activity analyses were performed using this TF activity assay.

### Differential expression and pathway enrichment analyses

Differential expression (DE) and functional pathway analyses, including over-representation analysis (ORA) and gene set enrichment analysis (GSEA), were performed on Seurat objects using SCP (v0.5.6; https://zhanghao-njmu.github.io/SCP) together with clusterProfiler (v4.10.1) and org.Hs.eg.db (v3.18.0)^58^. DE genes between conditions were identified with RunDEtest (SCP), which uses Seurat’s differential expression framework, with fc.threshold = 1 and only.pos = FALSE to capture both up- and downregulated genes. DE results were visualized using VolcanoPlot. ORA was conducted with RunEnrichment (SCP) against GO Biological Process (GO_BP) and KEGG gene sets for Homo sapiens, using DE genes filtered by avg_log2FC > log2(1) and p_val_adj < 0.05, and enriched terms were displayed using EnrichmentPlot. GSEA was performed with RunGSEA (SCP) using GO_BP gene sets for Homo sapiens and DE results filtered by p_val_adj < 0.05. Enrichment patterns were summarized with GSEAPlot in comparison mode, showing both positively and negatively enriched pathways.

### Cell-cell communication analysis

Intercellular communication networks were inferred using CellChat (v2.1.2)^26^, which predicts ligand–receptor–mediated signaling between cell populations and summarizes signaling inputs/outputs using network analysis and pattern-recognition frameworks. Gene expression matrices were extracted (RNA assay in Seurat objects) and used to construct CellChat objects. Overexpressed signaling genes and ligand–receptor interactions were identified using identifyOverExpressedGenes and identifyOverExpressedInteractions, and expression values were projected onto a protein–protein interaction (PPI) network. Communication probabilities were then estimated at the interaction and pathway levels using computeCommunProb and computeCommunProbPathway. Predicted signaling roles and aggregated communication networks were visualized using netVisual_signalingRole and netVisual_aggregate.

### Weighted gene co-expression network analysis

Gene co-expression networks were constructed using hdWGCNA (v0.4.00) on SCTransform-normalized single-cell RNA-seq data stored in a Seurat object^61^. SCTransform-scaled expression values were extracted from the SCT assay (GetAssayData, assay = “SCT”, slot = “scale.data”), and only genes retained by SCTransform (i.e., SCT features) were used as input for network construction. The Seurat object was prepared for hdWGCNA using SetupForWGCNA (wgcna_name = “SCT”, features = SCT features). For network inference, the expression matrix was set using SetDatExpr for a specified set of dopaminergic subtypes with. A soft-thresholding power was selected by evaluating scale-free topology and mean connectivity across candidate powers using TestSoftPowers and PlotSoftPowers; the final network was constructed at soft power = 10. Co-expression modules were identified using ConstructNetwork. Module eigengenes were computed with ModuleEigengenes, and intramodular connectivity metrics were calculated using ModuleConnectivity. Module assignments were retrieved with GetModules, excluding the unassigned “grey” module, and hub genes were defined as the top genes ranked by module membership (kME), using GetHubGenes (n_hubs = 20 unless otherwise indicated). Module structure and membership were visualized using dendrogram and kME plots (PlotDendrogram, PlotKMEs).

Module-level expression programs were quantified in single cells using ModuleExprScore with UCell scoring (method = “UCell”), and module eigengenes and/or module scores were visualized across embeddings and cell annotations using ModuleFeaturePlot (features = “hMEs” or “scores”) and Seurat-based summary plots (e.g., DotPlot grouped by “Celltype_final”). For presentation consistency, module names and colors were optionally reset using ResetModuleNames (new_name = “Module”) and ResetModuleColors (custom palettes), and harmonized module eigengenes were extracted with GetMEs (harmonized = TRUE) and appended to Seurat metadata for downstream visualization and statistical comparisons (e.g., FeatureStatPlot).

Module network representations were generated using ModuleNetworkPlot and HubGeneNetworkPlot (default parameters). Hub-gene embeddings were computed using RunModuleUMAP (n_hubs = 10, n_neighbors = 15, min_dist = 0.1) and visualized using ModuleUMAPPlot; module correlograms were generated using ModuleCorrelogram.

Functional enrichment analysis of module gene sets was performed using the Enrichr interface (enrichR package, v3.2) implemented in hdWGCNA (RunEnrichr). Enrichment was tested against Gene Ontology Biological Process, Cellular Component, Molecular Function (2021 releases) and KEGG (2021) databases, using up to 100 genes per module ranked by module membership (max_genes = 100; excluding the grey module). Enrichment results were retrieved using GetEnrichrTable and visualized using EnrichrBarPlot and EnrichrDotPlot with the number of displayed terms as indicated.

### Lineage inference and pseudotime-dependent gene dynamics

Trajectory inference was performed using Slingshot (v 2.10.0) and pseudotime-dependent gene expression dynamics were modeled using tradeSeq (v1.20.0)^39,40^. UMAP coordinates were extracted from the Seurat object and used as the reduced-dimensional input for lineage identification. Cell-type annotations were used as cluster labels. Slingshot lineages were inferred with getLineages on the UMAP embedding, specifying “progenitor” as the starting cluster (start.clus = “ progenitor”) and constraining terminal states to CALB1+_early_DA and ALDH1A1+_early_DA (end.clus = c(“CALB1+_early_DA”,“ALDH1A1+_early_DA”)). Principal curves were then fitted to each lineage using getCurves, and inferred lineages were visualized by overlaying curve trajectories on the UMAP projection.

For differential gene expression along pseudotime, a gene-by-cell count matrix was extracted from the Seurat RNA assay and restricted to genes used as SCTransform variable features (SCT var.features) to reduce noise and computational burden. The optimal number of knots for generalized additive model (GAM) fitting was assessed using evaluateK across a range of candidate values (k = 3–12; nGenes = 200). Slingshot pseudotime values and lineage assignment weights were obtained using slingPseudotime (na = FALSE) and slingCurveWeights, respectively. Gene expression dynamics were modeled by fitting lineage-aware GAMs using fitGAM with nknots = 10, using Slingshot pseudotime and cell weights as inputs. Global associations between gene expression and pseudotime were assessed using associationTest, and genes exhibiting distinct expression patterns across lineages were identified using patternTest. Significant genes were defined using a nominal p-value threshold (p ≤ 0.05 unless otherwise stated), and representative gene expression trends were visualized by plotting fitted smoothers along pseudotime using plotSmoothers.

### In silico gene perturbation analysis

In silico gene knockout (KO) analyses were performed using scTenifoldKnk (v1.3) on raw UMI count matrices derived from Seurat objects^62^. Briefly, cells of interest (e.g., A9 and A10 populations) were extracted from the filtered Seurat object, and a gene-by-cell count matrix was retrieved from the RNA assay (GetAssayData, slot = “counts”). Top 3,000 variable genes were retained and used as input for scTenifoldKnk.

Gene perturbation was simulated by specifying the KO gene (gKO; e.g., CREB5/ID4) in scTenifoldKnk. Quality control was enabled (qc = TRUE), with mitochondrial content filtering (qc_mtThreshold = 0.1) and a minimum library size threshold (qc_minLSize = 1000). For network construction, scTenifoldKnk was run using multiple subsampled cell sets (nc_nNet = 10) with nc_nCells = 500 cells per network and nc_nComp = 3 latent components; network regularization was set to nc_lambda = 0. Networks were scaled (nc_scaleScores = TRUE) without enforcing symmetry (nc_symmetric = FALSE), and edge selection was controlled by the quantile threshold nc_q = 0.9. Differential network analysis was performed using tensor decomposition with td_K = 3, a maximum of 1,000 iterations (td_maxIter = 1000), and convergence criteria td_maxError = 1e−5 (td_nDecimal = 3). Manifold/alignment steps were carried out with ma_nDim = 2. The resulting KO-associated network perturbation scores were used to prioritize downstream affected genes and pathways as indicated in subsequent analyses.

### Bulk RNA-seq differential expression analysis

Bulk RNA-seq read count matrices were analyzed using DESeq2 (v1.46.0)^80^. Gene counts were imported from the expression matrix and restricted to integer counts; genes with zero counts across all samples were removed. Sample metadata were parsed from sample identifiers to define dopaminergic subtype (A9 or A10), treatment (CTL or IFN-γ), and biological replicate (1 and 2). A combined group label (Subtype_Treatment) was used for summary and visualization. Replicate was included as a batch covariate where indicated.

Differential expression was modeled using a two-factor interaction design to test treatment effects within each subtype and to quantify subtype-specific treatment responses: ∼ Subtype * Treatment. Factor reference levels were set to A9 for subtype and CTL for treatment to facilitate interpretation of model coefficients. Low-count genes were optionally filtered prior to model fitting (retaining genes with total counts ≥ 10 across all samples). DESeq2 size factors and dispersions were estimated and negative binomial generalized linear models were fit using DESeq.

For visualization, variance-stabilized expression values were computed using the variance-stabilizing transformation (vst, blind = FALSE) and principal component analysis was performed using plotPCA, grouping samples by subtype and treatment. Differential expression contrasts were extracted as follows: the IFN-γ effect within A9 was obtained from the main treatment coefficient (Treatment_IFNg_vs_CTL); the interaction term (SubtypeA10.TreatmentIFNg) captured the difference in IFN-γ response between A10 and A9; and the IFN-γ effect within A10 was computed as the sum of the main treatment effect and the interaction term (contrast list combining Treatment_IFNg_vs_CTL and SubtypeA10.TreatmentIFNg). Genes were considered differentially expressed based on adjusted P values (Benjamini–Hochberg) and fold-change thresholds as specified in the corresponding figure legends.

### Multiomic analysis of chromatin accessibility and transcriptome

Single-cell Multiome-seq (ATAC+RNA) was performed on day 25 populations using the 10x Genomics Chromium Single Cell Multiome ATAC + Gene Expression platform at the Single Cell Innovation Lab (Memorial Sloan Kettering Cancer Center). Populations were individually labeled with TotalSeq hashtag oligonucleotides prior to pooling, with the following assignments: A0251, Day25_Control; A0252, Day25_ActA+BMP4; and A0253, Day25_ActA+BMP7. Hashtag-based demultiplexing was performed using Seurat’s HTODemux workflow as described above.

The data were processed in R (v4.4.2) using ArchR (v1.0.2) following the ArchR workflow (https://www.archrproject.com/bookdown/creating-an-archrproject-1.html)^44^. Fragment files generated by Cell Ranger ATAC (10x Genomics) were imported into ArchR using createArrowFiles with default parameters unless otherwise indicated.

A tile matrix was generated at 500-bp resolution using addTileMatrix, followed by iterative latent semantic indexing (LSI) with addIterativeLSI. Cell clustering was performed using addClusters and visualized with UMAP using addUMAP. Gene activity (gene score) matrices were computed using addGeneScoreMatrix (gene model: default ArchR annotations), and marker features were identified using getMarkerFeatures with bias correction for TSS enrichment and fragment counts (bias = c(“TSSEnrichment”, “log10(nFrags)”)). Motif annotations were added using addMotifAnnotations (motif set: CIS-BP/JASPAR, as specified), and deviations were calculated with chromVAR implemented in ArchR using addBgdPeaks and addDeviationsMatrix.

Peak calling was performed with MACS2 using pseudo-bulk replicates generated by ArchR (addGroupCoverages and addReproduciblePeakSet) followed by construction of a peak matrix with addPeakMatrix. Differential accessibility testing for peaks was carried out using getMarkerFeatures (testMethod = “wilcoxon”) with multiple hypothesis correction by false discovery rate (FDR). Peaks were linked to putative target genes using correlation-based peak-to-gene linkage (addPeak2GeneLinks), and browser tracks were generated with plotBrowserTrack from group-level coverage files. Unless otherwise indicated, statistical significance for differential features was defined as FDR < 0.05). To align chromatin accessibility with transcriptomic cell states, scATAC-seq clusters were integrated with matched scRNA-seq references using ArchR’s label transfer framework. Briefly, a Seurat object containing scRNA-seq data (day 25 annotated dataset) was used as a reference, and integration was performed with addGeneIntegrationMatrix using the gene score matrix from ArchR as the scATAC input and the normalized RNA expression as the reference. Predicted cell identities were assigned based on predictedGroup with associated confidence scores, and concordance between ATAC-derived clusters and RNA-derived annotations was assessed by comparing cluster-level label distributions. For visualization, integrated embeddings were displayed using ArchR UMAP coordinates and gene-level features were plotted using gene scores (scATAC) and normalized expression (scRNA) as indicated. Candidate positive transcriptional regulators were prioritized by integrating TF motif accessibility variability with correlation to TF activity. Motif accessibility deviations (chromVAR deviation Z-scores) were summarized at the cluster level using getGroupSE on the ArchR MotifMatrix (groupBy = “Clusters2”), and the Z-score component was extracted by subsetting rows with seqnames == “z”. For each motif, inter-cluster variability was quantified as the maximum pairwise difference in deviation Z-score (maxDelta) across clusters.

To identify TFs whose motif accessibility tracked with TF activity, motif deviation Z-scores were correlated with either gene activity (GeneScoreMatrix) or RNA-based TF expression (GeneIntegrationMatrix) using correlateMatrices across low-overlap cell aggregates in the IterativeLSI space. The resulting correlation tables were annotated with motif maxDelta values. Positive TF regulators were defined as TFs with motif–activity correlation > 0.5, adjusted P < 0.01, and maxDelta in the top quartile, after collapsing redundant motif entries to unique TF names. Associations were visualized by plotting correlation versus maxDelta and highlighting TFs meeting these criteria.

### Electrophysiological recordings in cultured mDA neurons

Patch-clamp electrophysiological recordings were performed using previous described method^8^. Specifically, the assay was performed on hPSC-derived mDA neurons at day 60 at room temperature in a Tyrode’s solution containing (in mM): 119 NaCl, 3 KCl, 10 glucose, 2 CaCl2, 1.2 MgCl2-6 H2O, 3.3 HEPES, and 2.7 HEPES-Na+ salt (pH 7.4, 270 mm). For whole-cell patch-clamp studies, borosilicate glass pipettes (G150F-4, Warner Instruments, CT) with a tip resistance of 3-4 MΩ were pulled on a P-97 Flaming-Brown micropipette puller (Sutter Instruments, CA) and filled with (in mM): 115 K-gluconate, 20 KCl, 10 HEPES, 2 MgCl2, 2 ATP-Mg, 2 ATP-Na2 and 0.3 GTP-Na, (pH 7.25, ∼280 mOsm). Neurons were visualized under a 40x water immersion objective using Olympus BX51W1 microscope (Olympus), and recording were performed with an MultiClamp 700B amplifier (Molecular Devices, CA) and digitized at 10 kHz with ITC-18 (HEKA Instruments Inc, NY). Data were acquired using WinWCP software (John Dempster, University of Strathclyde, UK). In each cell, input resistance (measured by −100 pA, 1s hyperpolarizing pulse), resting membrane potential and spontaneous action potentials were monitored throughout the recording. Current-voltage relationship and evoked action potentials were measured by injecting a somatic current (1s duration) from −30 to +20 pA in +10 pA increments and from 0 to +250 pA in +10 pA increments, respectively.

### High-Performance Liquid Chromatography (HPLC)

HPLC was performed using the same method previously described^8^. mDA neurons were plated on PO/laminin/fibronectin coated 24-well plates at 5 × 10^5^ cells/well density on day 25 and used at day 60. HPLC with electrochemical detection was done as previously described^91^. Briefly, cells were preincubated in fresh DMEM: F12 + N2 media for 30 min. After exposure to either Tyrode’s saline alone or supplemented with high KCl (80 mM, Sigma) for 5 min at 37°C, supernatant was collected and immediately mixed with 0.2 M perchloric acid (1:1 volume) to deproteinize the sample and prevent dopamine auto-oxidation. Perchloric acid was also added into the wells with cells to measure intracellular DA concentration. After 10 min incubation at room temperature, samples were centrifuged at 10,000 g for 5 min at 4°C, supernatant was collected, stored at -80°C and analyzed within the following two weeks. DA concentrations in each group of samples were normalized to the levels in the corresponding control group; data were averaged from 2 independent experiments. Intracellular and extracellular DA levels were divided by the fraction of DA neurons in each group, obtained from immunostaining of sister cultures for TH.

### Cell transplantation, tissue processing and immunohistochemistry

All animal procedures were performed at Memorial Sloan Kettering Cancer Center in accordance with protocols approved by the Institutional Animal Care and Use Committee and in compliance with NIH guidelines. NOD.Cg-*Prkdc*^scid^ *Il2rg*^tm1Wjl^/SzJ mice (NSG; 6–8 weeks old; The Jackson Laboratory) were used for transplantation experiments. Mice were anesthetized with isoflurane throughout the surgical procedure. Cell aggregates were resuspended in transplantation medium consisting of Neurobasal medium supplemented with 2 mM L-glutamine, 0.2 mM ascorbic acid, 0.1% Kedbumin, 1 mg/mL TNFα inhibitor (adalimumab)^92^, and CEPT cocktail. A total of 40 cell aggregates (10,000 cells/aggregate) in 10 μl were stereotactically injected into the striatum using a motorized stereotaxic injector (Model 53311, Stoelting) at a rate of 1 μl/min. Cells were delivered across two injection tracks at the following coordinates relative to bregma: anteroposterior (AP) +0.3 mm, mediolateral (ML) −2.1 mm, dorsoventral (DV) −3.2, −3.1, −3.0 and −2.9 mm; and AP +1.0 mm, ML −1.7 mm, DV −3.2, −3.1, −3.0 and −2.9 mm. Following injection, the needle was left in place for 5 min and then slowly withdrawn at 1 mm/min.

At the indicated endpoints, animals were transcardially perfused with 4% paraformaldehyde (PFA). Brains were dissected, post-fixed in 4% PFA for 18 h, cryoprotected in 30% sucrose prepared in 0.01 M phosphate-buffered saline (PBS) for 24 h, embedded in O.C.T. Compound (Sakura Finetek), and cryosectioned at 20 μm thickness. For immunohistochemistry, tissue sections were washed three times in PBS and blocked for 1 h at room temperature in PBS containing 1% bovine serum albumin and 0.3% Triton X-100. Sections were incubated overnight at 4 °C with primary antibodies against GIRK2 (1:100; Alomone Labs, APC_006-GP) and calbindin (1:200; Cell Signaling Technology, 13176). After three washes in PBS, sections were incubated for 1 h at room temperature with appropriate Alexa Fluor 488- or 647-conjugated secondary antibodies (1:400; Thermo Fisher Scientific), followed by nuclear counterstaining with DAPI. Stained sections were scanned at the MSKCC Molecular Cytology Core Facility using a Pannoramic Scanner (3DHISTECH) equipped with a 20×/0.8 NA objective. Images were acquired and processed using CaseViewer software (v.2.4; 3DHISTECH)

### Quantification and statistical analysis

Standard statistical analyses were performed using GraphPad Prism 10. The number of replicates, type of statistical test and test results are described in the figure legends. All data are represented as mean ± s.e.m. Results are significant at p < 0.05 (∗), p < 0.01 (∗∗), p < 0.001 (∗∗∗), p < 0.0001 (∗∗∗∗). Sample size of all the experiments was not pre-determined, and no randomization or investigator blinding approaches were implemented during the experiments and data analyses given the nature of the study.

## Data availability

All raw and processed data generated from the bulk and single-cell sequencing studies reported in this manuscript have been deposited in the EMBL-EBI repository under accession numbers E-MTAB-17399, E-MTAB-17418, E-MTAB-17419 and E-MTAB-17424.

## Code availability

No custom software or algorithms were developed for this study. All computational analyses were performed using publicly available bioinformatics tools, as described in the Methods.

