## Supplemental figures for "Decoding subtype development and function in human pluripotent stem cell-derived midbrain dopaminergic neurons": Extended Figures and Captions biorxv 0923.pdf

### Extended Figure 1

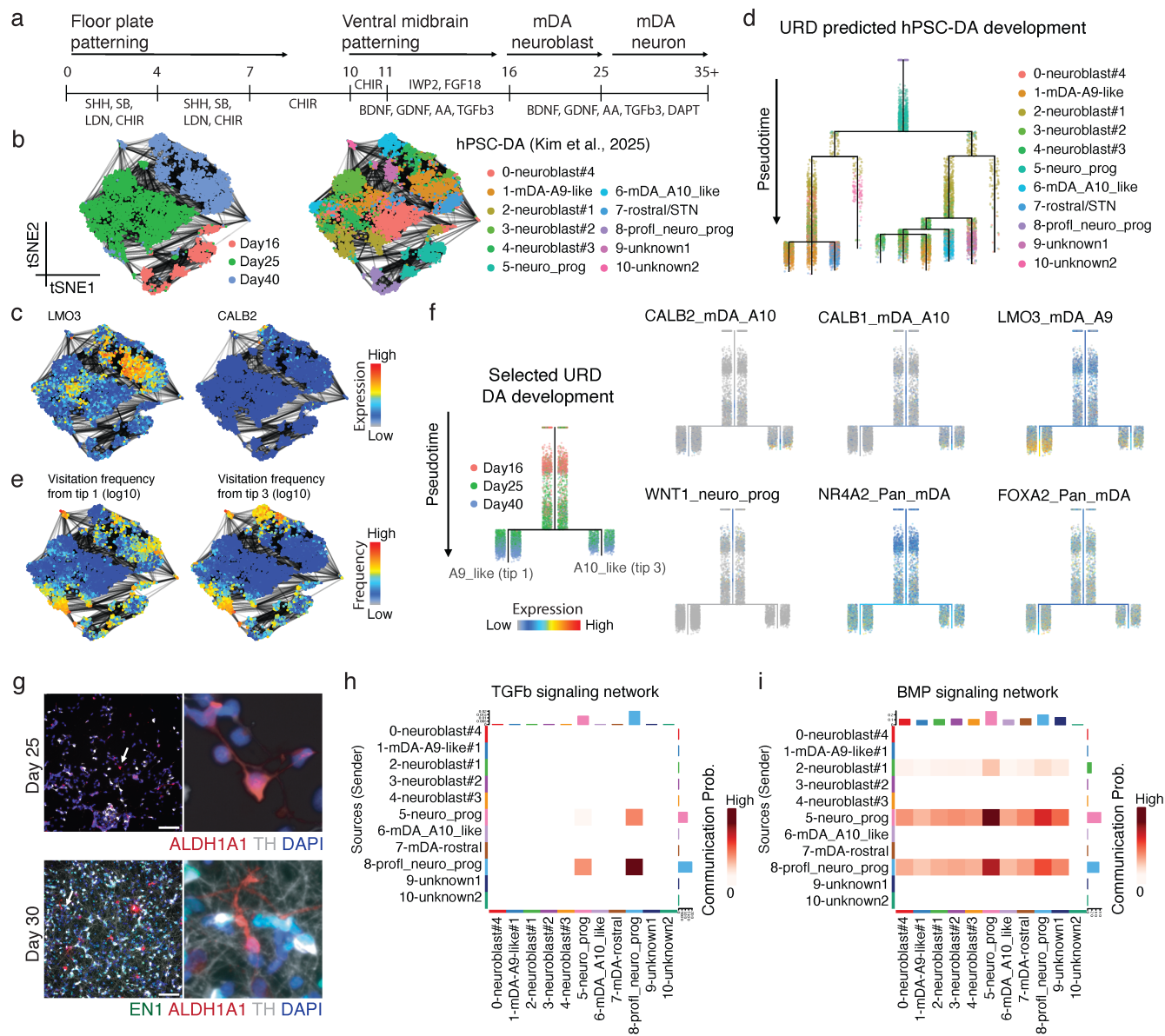

**Extended Fig. 1 | Identification of candidate time windows and regulators of mDA subtype specification.**

- **a**, Schematic of the hPSC-to-mDA neuron differentiation protocol and sampled time points for single-cell profiling.
- **b**, tSNE projection of hPSC-derived cells colored by differentiation stage (left) and cell-type annotation (right).
- **c**, tSNE feature plots showing expression of the A10-enriched markers *LMO3* and *CALB2*.
- **d**, URD pseudotime trajectory reconstruction of mDA differentiation.
- **e**, tSNE plots showing URD visitation frequency from terminal states (tips) 1 and 3.
- **f**, URD trajectories colored by differentiation stage (left) and expression of selected genes (right).
- **g**, Representative IF images showing ALDH1A1 and TH at day 25, and EN1, ALDH1A1 and TH at day 30. Scale bars, 100 $\mu\text{m}$.
- **h,i**, Network heatmaps summarizing inferred activity of TGFβ (h) and BMP (i) signaling programs across the indicated populations.

#### Extended Figure 2

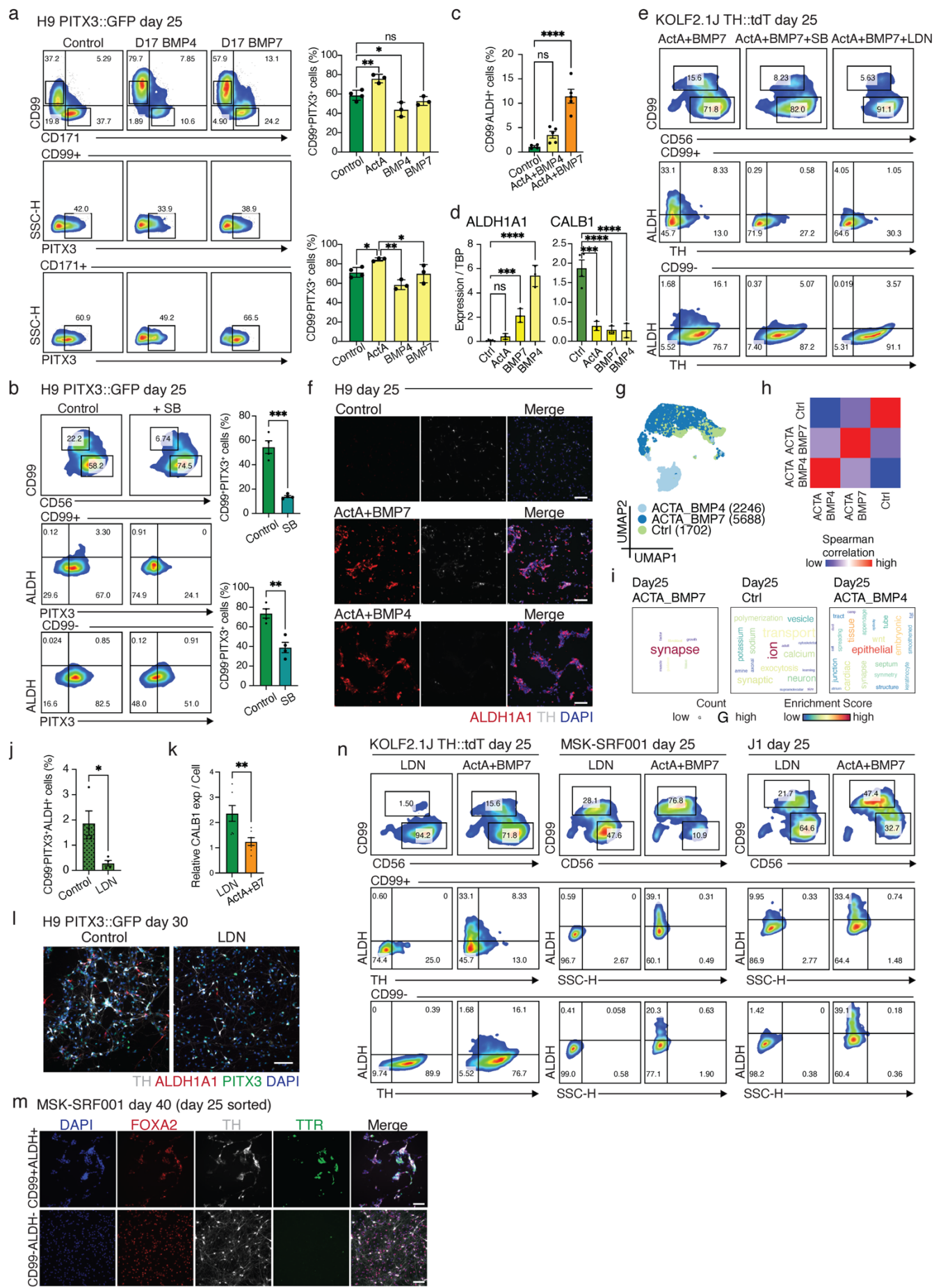

**Extended Fig. 2 | Related to Fig. 1.**

- **a**, Flow-cytometric quantification of PITX3, CD99 and CD171 across the indicated populations. One-way ANOVA; $n = 3$.
- **b**, Flow-cytometric quantification of PITX3 expression and ALDH activity in the indicated populations stratified by CD99 and CD56. Student's $t$-test; $n = 4$ independent differentiations.
- **c**, Quantification of the CD99–ALDH+ fraction (related to Fig. 1d). Student's $t$-test; $n = 4$-5 independent differentiations.
- **d**, RT–qPCR analysis of *ALDH1A1* and *CALB1* expression in the indicated populations. One-way ANOVA; $n = 3$.
- **e**, Flow cytometry of TH expression and ALDH activity in the indicated CD99– and CD99+ populations.
- **f**, Immunofluorescence analysis of ALDH1A1 and TH expression in day 25 populations generated under the indicated conditions. Scale bars, 100 $\mu\text{m}$.
- **g**, UMAP embedding of D25 populations.
- **h**, Spearman correlation of pseudo-bulk transcriptomes for the indicated populations.
- **i**, Pathway enrichment analysis based on genes upregulated in the indicated populations.
- **j**, Quantification of the D25 CD99–PITX3+ALDH+ fraction (related to Fig. 1g). Student's $t$-test; $n = 4$ independent differentiations.
- **k**, Quantification of relative CALB1 expression in D26 populations (related to Fig. 1h). Student's $t$-test; $n = 7$.
- **l**, Representative IF images showing TH, ALDH1A1 and PITX3 expression in D30 populations. Scale bars, 100 $\mu\text{m}$.
- **m**, Representative IF images showing TH, FOXA2 and TTR expression in the indicated D40 populations. Scale bars, 100 $\mu\text{m}$.
- **n**, Representative flow cytometry of TH, CD56 and CD99 expression and ALDH activity in D25 populations derived from KOLF2.1, MSK-SRF001 and J1 hPSC lines. All error bars in the figure represent s.e.m.

### Extended Figure 3

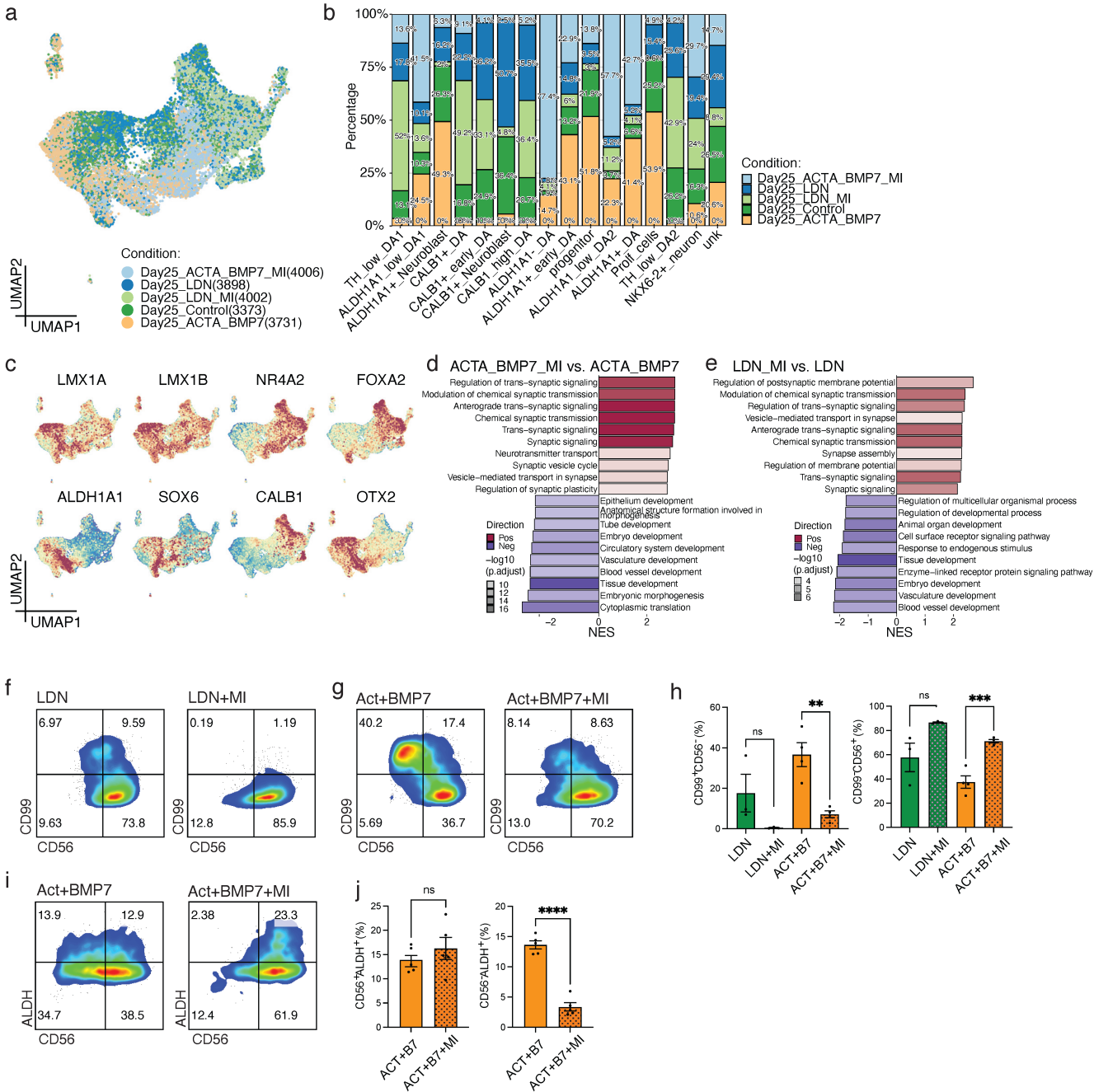

**Extended Fig. 3 | Related to Fig. 2.**

**a**, UMAP of D25 populations colored by experimental condition.

**b**, Bar plots showing the relative contribution of each condition to each annotated cell type.

**c**, UMAP feature plots showing expression of the indicated genes.

**d,e**, GSEA comparing control versus MEK inhibitor–treated populations patterned with Activin A and BMP7 (d) or LDN (e).

**f–h**, Flow cytometry of CD56 and CD99 expression in the indicated A10-patterned (f) and A9-patterned (g) populations, with quantification of CD99+ and CD99– fractions (h). Student's *t*-test; *n* = 4.

**i,j**, Flow cytometry of CD56 and ALDH activity in A9-patterned populations ± MEK inhibitor (i), with quantification of CD56+ALDH+ and CD56–ALDH+ fractions (j). Student's *t*-test; *n* = 4 independent differentiations; all error bars in the figure represent s.e.m.

Extended Figure 4

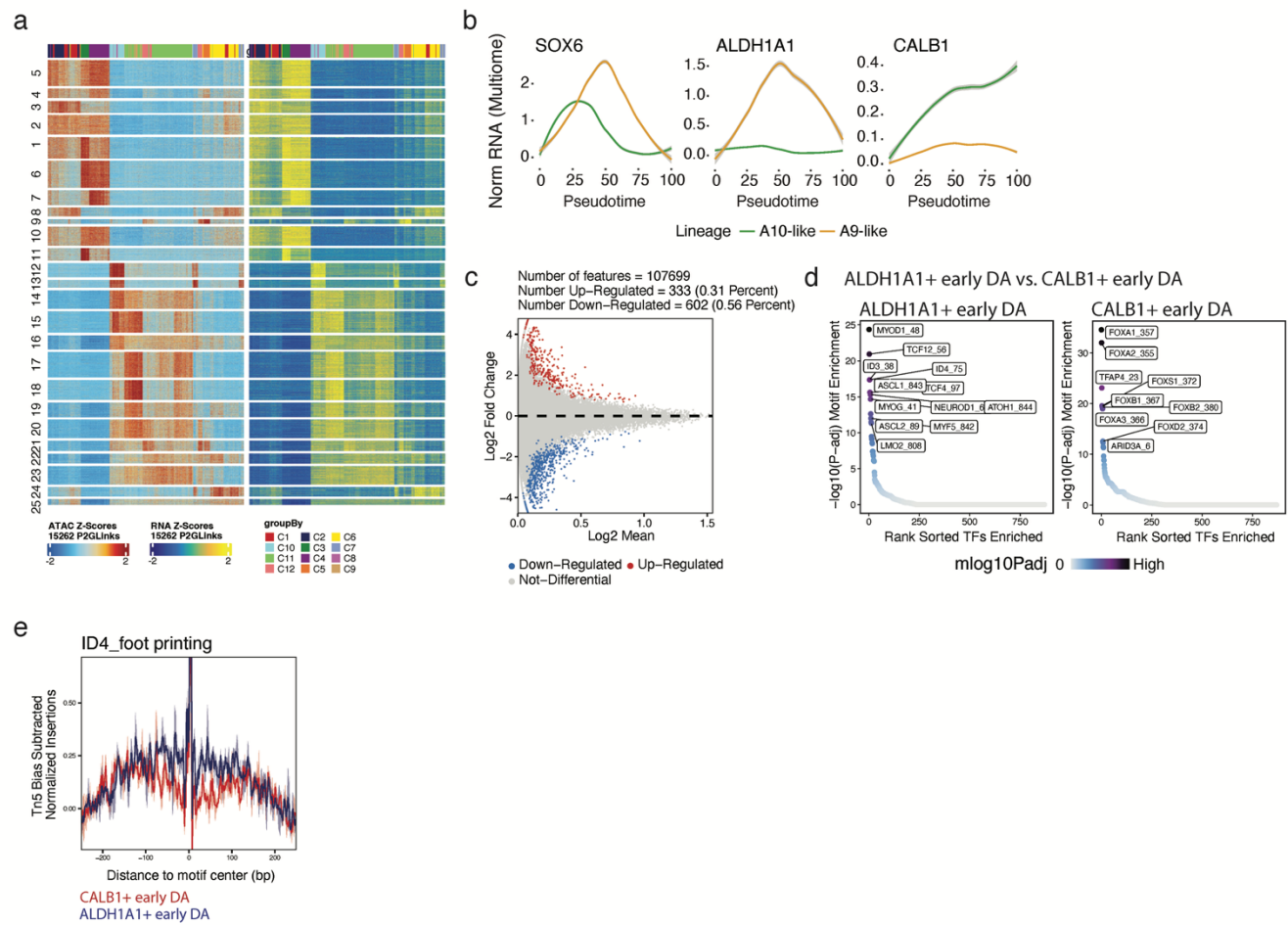

**Extended Fig. 4 | Related to Fig. 2.**

**a,** Heatmaps summarizing z-scored chromatin accessibility (ATAC) and gene expression (RNA) from single-cell Multiome profiling.

**b,** Expression of the indicated genes along A9-like and A10-like pseudotime trajectories.

**c,d,** Motif enrichment analysis comparing ALDH1A1+ early DA and CALB1+ early DA clusters.

**e,** Transcription factor footprinting analysis highlighting ID4 occupancy.

### Extended Figure 5

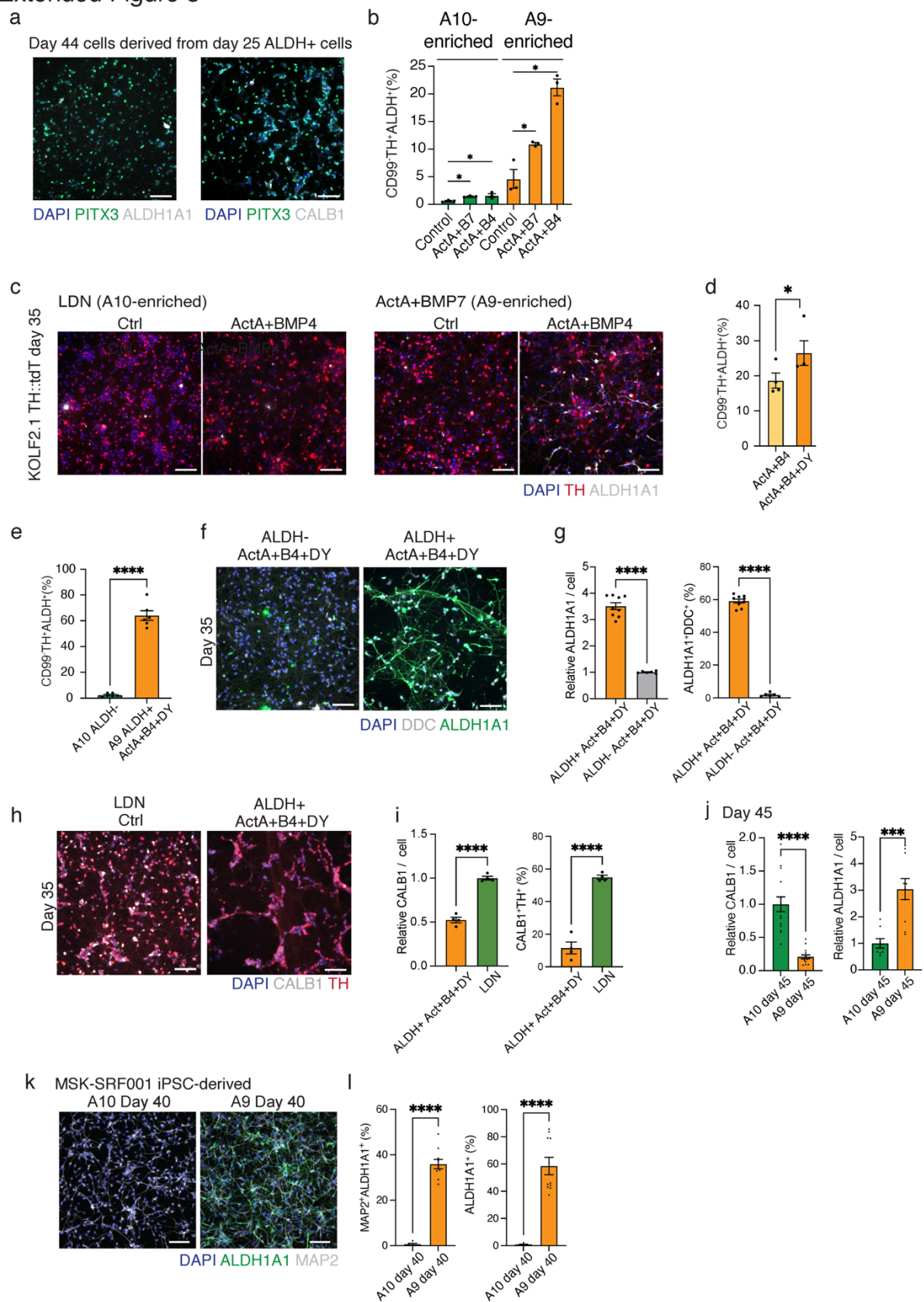

**Extended Fig. 5 | Related to Fig. 3.**

- **a**, Representative IF images showing PITX3 and ALDH1A1 expression, and PITX3 and CALB1 expression, in day 44 populations derived from day 25 ALDH+ A9 cells. Scale bars, 100 μm.
- **b**, Quantification of CD99-TH+ALDH+ cells in D35 A9- and A10-enriched populations treated with the indicated factors (related to Fig. 3c,d). Student's *t*-test; *n* = 3.
- **c**, IF images showing TH and ALDH1A1 in D35 A9- and A10-enriched populations generated using the indicated conditions. Scale bars, 100 μm.
- **d**, Quantification of CD99-TH+ALDH+ cells in D35 A9-enriched populations treated with the indicated factors (related to Fig. 3e). Student's *t*-test; *n* = 4.
- **e**, Quantification of CD99-TH+ALDH+ cells in day 35 A10 and A9 populations treated with A9-maintenance factors (related to Fig. 3h). Student's *t*-test; *n* = 6.
- **f,g**, Representative IF images of DDC and ALDH1A1 in D35 populations derived from D25 sorted ALDH- and ALDH+ cells and cultured under A9-maintenance conditions (f). Quantification of relative ALDH1A1 expression and the proportion of ALDH1A1+DDC+ cells in D35 populations (g). Student's *t*-test; *n* = 6–9; error bars, s.e.m.
- **h,i**, Representative IF images of CALB1 and TH in D35 populations derived from D25 sorted ALDH- (LDN/A10-patterned) and ALDH+ cells and cultured under control or A9-maintenance conditions (h). Quantification of relative CALB1 expression (left) and CALB1+TH+ proportions (right) in D35 populations (i). Student's *t*-test; *n* = 4.
- **j**, Quantification of relative CALB1 and ALDH1A1 expression in D45 populations (related to Fig. 3i). Student's *t*-test; *n* = 8.
- **k,l**, Representative IF images of ALDH1A1 and MAP2 in day 40 A10 and A9 neurons derived from MSK-SRF001 iPSCs (k), and quantification of ALDH1A1 expression and the proportion of ALDH1A1+ MAP2+ cells in day 40 mDA subtypes derived from MSK-SRF001 iPSCs (l). Student's *t*-test; *n* = 7; all error bars in the figure represent s.e.m.

#### Extended Figure 6

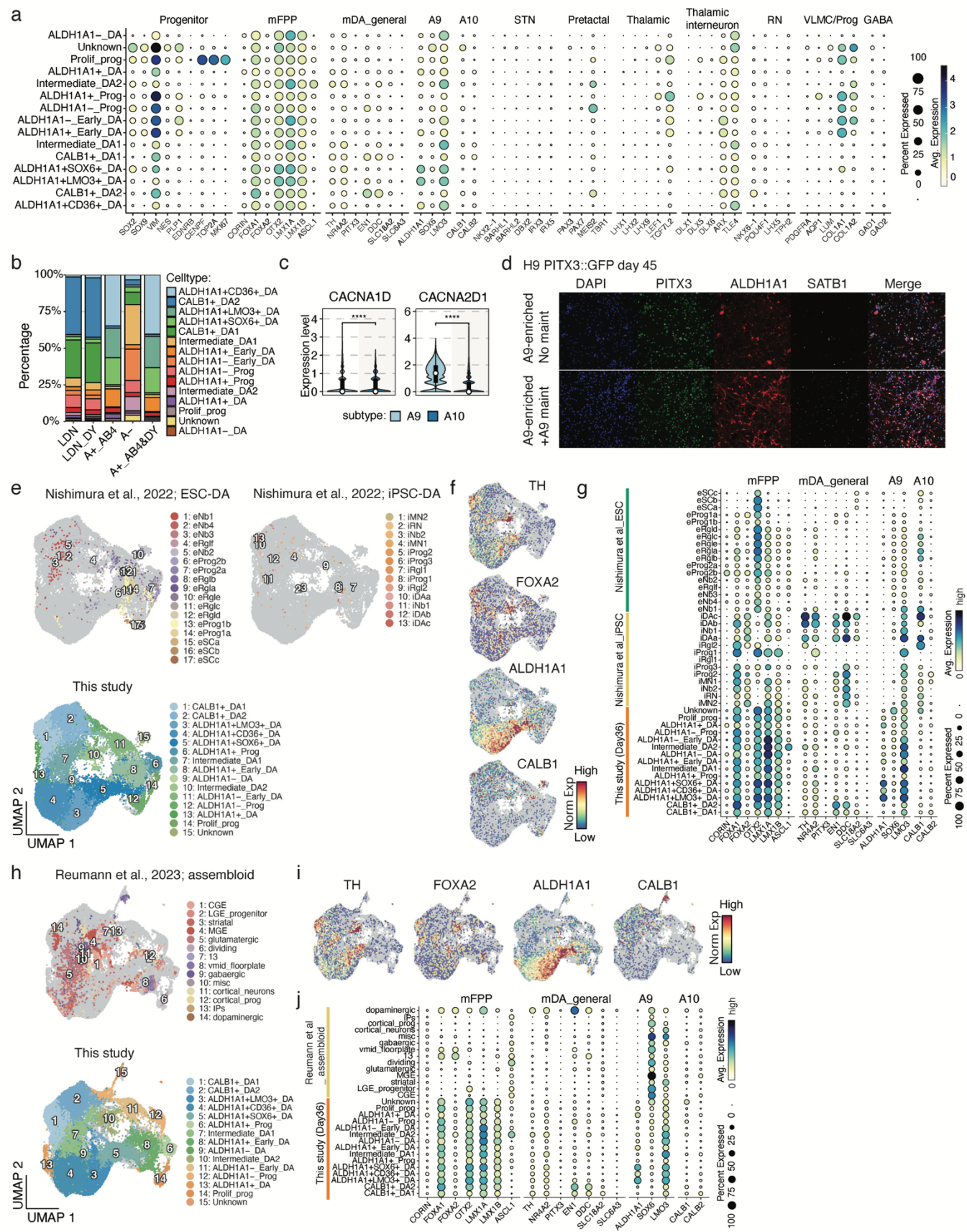

**Extended Fig. 6 | Related to Fig. 4.**

- **a**, Dot plot showing expression of gene categories across annotated cell types (related to Fig. 4a).
- **b**, Bar plot showing the relative contribution of cells from each differentiation condition to each annotated cell type (related to Fig. 4a).
- **c**, Expression of *CACNA1D* and *CACNA2D1* in pseudo-bulk A9 and A10 groups. Wilcoxon test.
- **d**, Representative IF images showing PITX3, ALDH1A1 and SATB1 in D45 A9-patterned populations cultured with or without A9 maintenance conditions.
- **e**, CCA-based integration of D36 hPSC-mDA data (this study) with published ESC- and iPSC-derived mDA datasets.
- **f**, UMAP feature plots showing expression of the indicated genes in the integrated dataset.
- **g**, Dot plot comparing expression of the indicated genes in the integrated dataset, stratified by dataset of origin.
- **h**, Integration of D36 hPSC-mDA data (this study) with published midbrain assembloid datasets.
- **i**, UMAP feature plots showing expression of the indicated genes in the integrated dataset.
- **j**, Dot plot comparing expression of the indicated genes in the integrated dataset, separated by dataset of origin.

Extended Figure 7

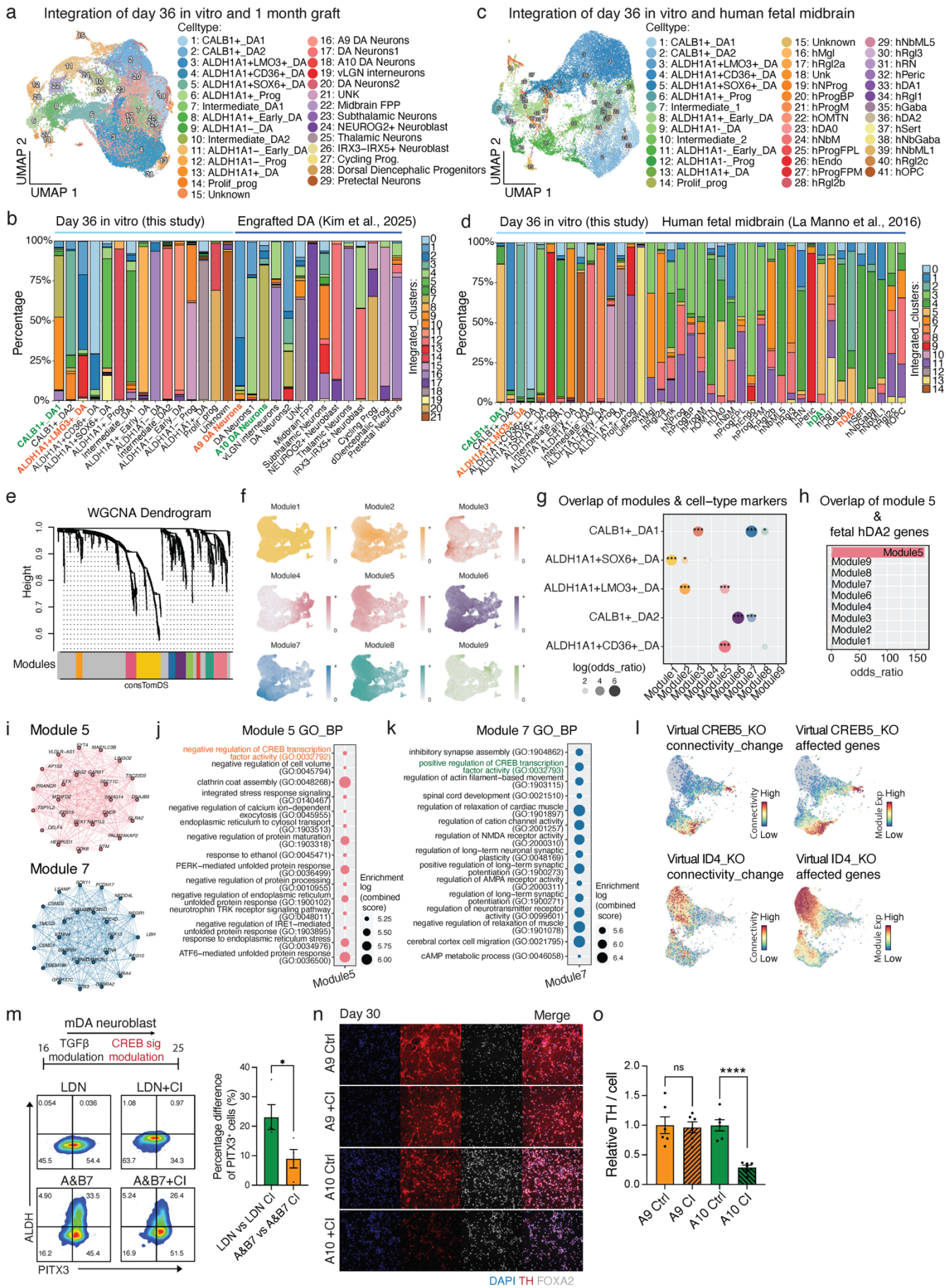

**Extended Fig. 7 | Related to Fig. 4.**

- **a**, CCA-based integration of D36 hPSC-mDA data (this study) with grafted hPSC-DA data (related to Fig. 4j).
- **b**, Bar plot showing the relative contribution of annotated cell types from each data source to integrated clusters (related to Fig. 4j).
- **c**, CCA-based integration of D36 hPSC-mDA data (this study) with human fetal midbrain data (related to Fig. 4o).
- **d**, Bar plot showing the relative contribution of annotated cell types from each data source to integrated clusters (related to Fig. 4o).
- **e,f**, hdWGCNA module dendrogram (e) and UMAP plots showing gene-module expression across cells (f).
- **g**, Overlap between hdWGCNA modules and genes upregulated in the indicated cell types (this study).
- **h**, Overlap between hdWGCNA modules and genes upregulated in the hDA2 cluster in human fetal midbrain data.
- **i-k**, Representative genes within modules 5 and 7 (i) and pathway enrichments for module 5 (j) and module 7 (k).
- **l**, UMAP showing network connectivity changes and expression patterns of dysregulated genes following in silico knockout of *CREB5* (top) or *ID4* (bottom).
- **m**, Flow-cytometric analysis of ALDH activity and PITX3 expression in the indicated populations ± CREB inhibitor (left), with quantification of PITX3+ cells (right). Student's *t*-test; *n* = 3 differentiations.
- **n,o**, Representative IF images of D30 A9 and A10 populations generated under the indicated conditions (n), with quantification of relative TH expression (o). Student's *t*-test; *n* = 6; all error bars, s.e.m.

#### Extended Figure 8

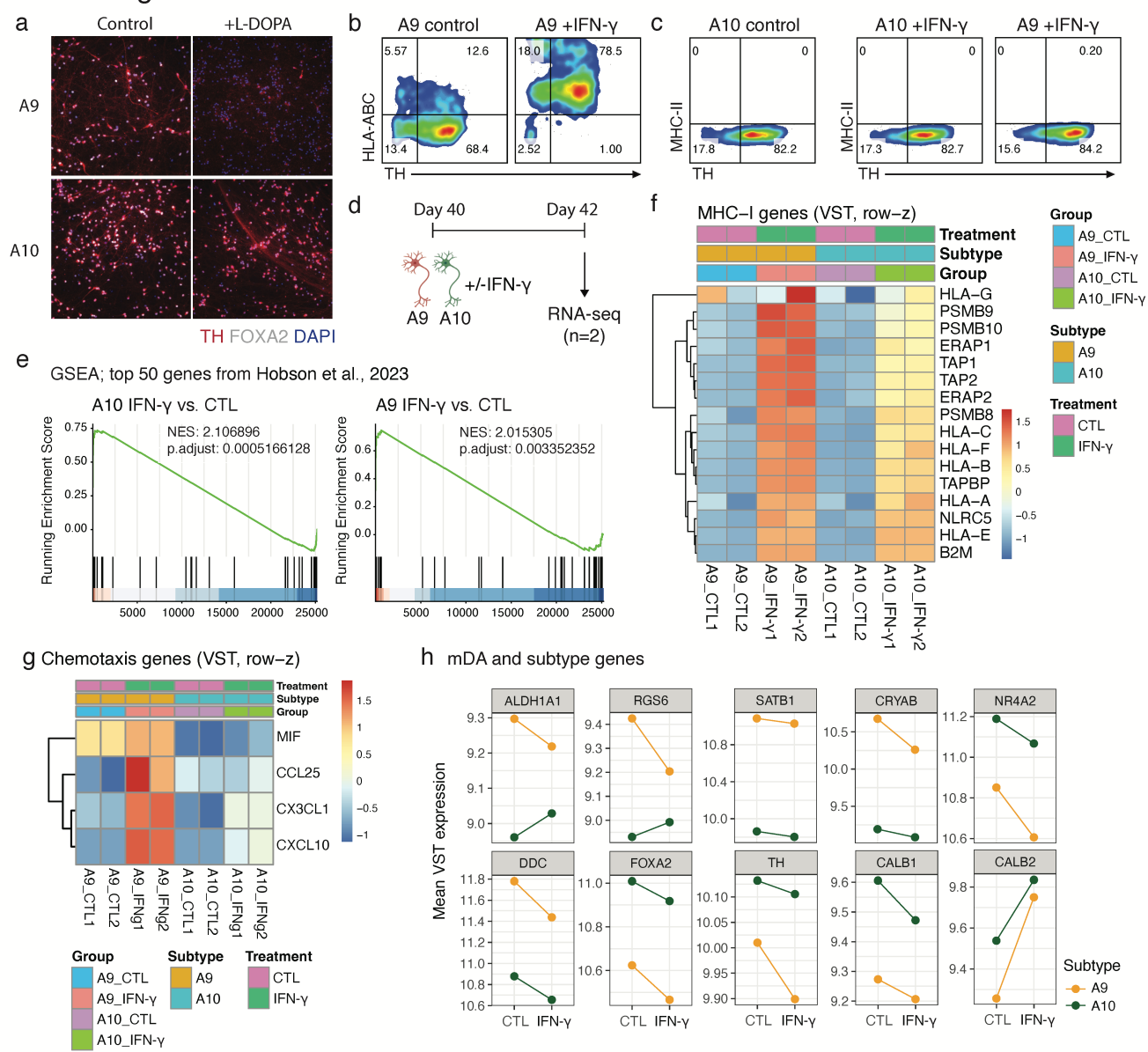

**Extended Fig. 8 | Related to Fig. 6.**

- **a**, Representative IF images of FOXA2 and TH in D50 A9 and A10 cultures ± L-DOPA.
- **b**, Representative flow-cytometric analysis of TH and HLA-ABC in A9 cultures ± IFN-γ.
- **c**, Representative flow-cytometric analysis of TH and MHC-II in A9 and A10 cultures ± IFN-γ.
- **d**, Schematic of bulk RNA-seq experimental design for A9 and A10 cultures ± IFN-γ.
- **e**, GSEA of mouse IFN-γ-responsive gene sets in IFN-γ-treated hPSC-A9 and hPSC-A10 relative to matched controls.
- **f–h**, Variance-stabilized (VST) expression of MHC-I genes (f), chemotaxis-associated genes (g), and mDA/subtype marker genes (h) in A9 and A10 cultures ± IFN-γ.
